# Dopaminergic neurons dynamically update sensory values during navigation

**DOI:** 10.1101/2022.08.17.504092

**Authors:** Ayaka Kato, Kazumi Ohta, Kazuo Okanoya, Hokto Kazama

## Abstract

Updating the value of sensory cues through experiences is a critical element of adaptive behavior. Although dopaminergic neurons (DANs) achieve this by driving associative learning, whether they contribute to assessment of sensory values outside the context of association remains largely unexplored. Here we show in *Drosophila* that DANs in the mushroom body encode the innate value of odors and constantly update the current value by inducing plasticity during olfactory navigation. Simulation of neuronal activity in a network based on the connectome data reproduced the characteristics of DAN responses, proposing a concrete circuit mechanism for computation. Notably, odors alone induced value- and dopamine-dependent changes in the activity of mushroom body output neurons, which store the current value of odors, as well as the behavior of flies steering in a virtual environment. Thus, the DAN circuit known for discrete, associative learning also continuously updates odor values during navigation in a non-associative manner.

## Introduction

Animals explore the environment based on the value of sensory cues, which is constantly updated through experiences. Updating of sensory values has been most extensively studied in the context of associative learning, where sensory cues acquire new values from reinforcers such as reward and punishment (Pavlov, 1927; Rescorla, 1972; Thorndike, 1911). Dopaminergic neurons (DANs) have been shown to play a central role in this process by encoding information about reinforcers (Schultz et al., 1997; Spanagel and Weiss, 1999; Watabe-Uchida et al., 2017), and inducing plasticity at sensory synapses (Calabresi et al., 2007; Tritsch and Sabatini, 2012). This plasticity requires coincident recruitment of dopaminergic and sensory pathways within a narrow time window (Black et al., 1985; Smith-Roe and Kelley, 2000; Yagishita et al., 2014).

Although DAN’s responses are thoroughly characterized during association where sensory cues and reinforcers are given repeatedly in combination, DANs also respond to natural stimuli such as food and sweet taste presented in isolation (Bassareo and Di Chiara, 1999; Hajnal et al., 2004), suggesting that DANs encode innate value of stimuli in general. Given that a variety of sensory stimuli including odors that are typically used as conditioned stimuli possess innate values (Ohman and Mineka, 2001; Richardson and Zucco, 1989; Yarmolinsky et al., 2009), certain stimuli alone may activate DANs simultaneously with sensory pathways and fulfill the very criteria for inducing synaptic plasticity without the additional reinforcement. If this were the case, sensory values would be updated continuously in a dopamine-dependent manner as the animal navigates the environment, a hypothesis that has not received scrutiny.

Here we examine this hypothesis in *Drosophila* navigating the olfactory environment by capitalizing on the following advantages that flies offer. First, flies exhibit inherent, graded responses to odors ranging from attraction to aversion, allowing us to infer the innate value of odors for the animals (Badel et al., 2016; Knaden et al., 2012; Semmelhack and Wang, 2009). Second, the anatomical organization of the olfactory circuit is well suited to comprehensively record the activity of related DANs. The olfactory association center, the mushroom body (MB), is subdivided into 15 compartments (Figure 1A), each of which houses a circuit comprising Kenyon cells (KCs) that convey olfactory information, MB output neurons (MBONs) that drive attraction or aversion, and DANs that modulate the synaptic strength between KCs and MBONs during olfactory association (Aso and Rubin, 2016; Aso et al., 2014a, 2014b; Endo et al., 2020; Hige et al., 2015; Owald et al., 2015; Turner et al., 2008). Each type of DAN innervates a single or a few compartments. This compact and segregated organization allows physiological access to the entire population of DANs in the MB.

**Figure 1.**
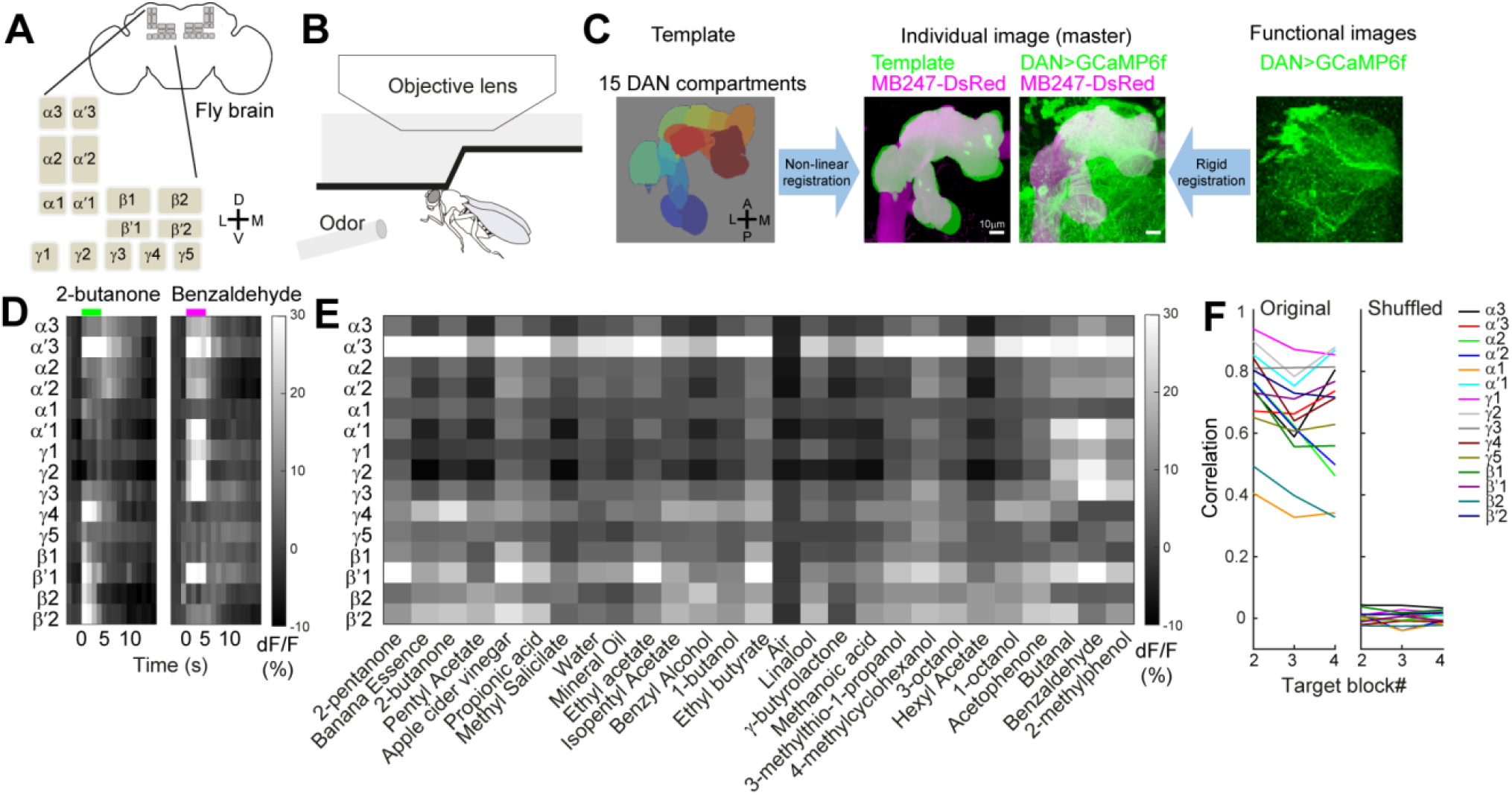
DANs in the MB exhibit stereotypical responses to a large panel of odors (A) Schematic of the fly brain and the 15 MB compartments. (B) Setup for volumetric Ca^2+^ imaging of odor-evoked DAN responses. (C) Schematic of the image registration process. A template volume of the 15 MB compartments (JFRC 2013; Aso et al., 2014a) was registered to MB247-DsRed signals (magenta) in a master image of individual brains co-expressing GCaMP6f (green) under the control of *TH-Gal4* and *Ddc-*Gal4 (Scale bar, 10 µm). Functional images were registered to the master image using GCaMP6f signals. (D) Average responses of DANs to representative attractive (2-butanone) and aversive (benzaldehyde) odors shown as ΔF/F of GCaMP6f fluorescence. Average is taken across trials and flies. Colored bars at the top indicate the period of odor application (4 s). (E) Average peak responses of DANs to 27 odors. Average is taken across time (odor application period), trials, and flies. Odors are arranged by innate values (attractive odors on the left). (F) Consistency of DAN responses across blocks in individual compartments quantified with Pearson’s correlation of peak responses to the 27 odors between block#1 and other blocks, with (right) or without (left) shuffling the odor identity in each block. Correlations for the original dataset were not significantly different across blocks (p = 0.16, Kruskal-Wallis test) but were higher than those for the shuffled dataset (p = 6.4×10^−16^, Friedman’s test). Data in D-F are from the same 10–11 flies. The number of flies varies between odors.

As in other organisms, DANs in flies have been known for their salient responses to reward and punishment (Kirkhart and Scott, 2015; Liu et al., 2012; Riemensperger et al., 2005; Tomchik, 2013), but they also inherently respond to odors (Dylla et al., 2017; Hattori et al., 2017; Mao and Davis, 2009; McCurdy et al., 2021; Riemensperger et al., 2005; Siju et al., 2020), with one study showing that DANs encode binary odor valence (Siju et al., 2020). Moreover, increasing reports suggest that odor-induced dopamine release exerts an effect on learning. For example, odor-induced DAN activity contributes to familiarization of stimuli (Hattori et al., 2017), extinction as well as reconsolidation of memory (Felsenberg et al., 2017, 2018), and latent inhibition (Jacob et al., 2021). However, how the population of DANs respond to odors alone in a more natural context (i.e. without reinforcers) and the extent to which this can allow dynamic updating of odor values remain unknown.

In this study, we investigated the olfactory computation performed by DANs, its underlying mechanism, as well as its physiological and behavioral consequences by combining two-photon functional imaging, connectome-based simulation, and a closed-loop behavioral analysis. We recorded the activity of DANs in all the MB compartments and found that DANs encode innate value of odors, ranking them from strongly attractive to strongly aversive. Simulation using a neural network model based on the connectome data (Scheffer et al., 2020) recapitulated the characteristics of DAN odor responses, suggesting the circuit mechanism underlying DAN activity. Notably, repetitive application of odors alone induced plastic changes in MBONs in a dopamine and odor value-dependent manner. Finally, optogenetic experiments showed that DANs drive odor value-dependent flight control in accordance with the plasticity of MBONs. Together, these results reveal that DANs recognized in associative learning are also active during navigation to continuously update odor values.

## Results

### DANs in the MB exhibit stereotypical responses to a large panel of odors

We characterized odor-evoked responses of DAN axons in the entire MB lobes encompassing all the 15 compartments using two-photon calcium imaging (Figures 1A-C). *TH-GAL4* and *Ddc-GAL4* were used in combination to express the calcium sensor GCaMP6f throughout DANs in the MB, and MB247-DsRed was used to delineate the MB lobes (Cohn et al., 2015; Riemensperger et al., 2005). To quantify the change in fluorescence in individual compartments, we non-linearly registered the three-dimensional template of the compartments (JFRC2013 standard brain, (Aso et al., 2014a)) to the DsRed signals in a high-resolution, master image (Figures 1C, S1). In parallel, functional images capturing odor responses were registered to the master image using the GCaMP6f signals (Figure 1C). We aligned these images to extract the activity of DANs in the 15 compartments.

We recorded responses of DANs to 27 different odors with a wide range of values (termed here as value index), previously quantified based on the behavioral responses of flies to these odors in a flight arena (Badel et al., 2016). DAN axons in different compartments exhibited distinct tuning and temporal responses to odors (Figure 1D, E). The tuning of peak responses remained stable across repeated application of odors (Figure 1F). The trial-to-trial variability of responses was similar between DANs and projection neurons (PNs), the second-order olfactory neurons in the antennal lobe (Figure S2). DAN’s odor responses mostly originate in the olfactory organs as they were largely diminished by removal of the antennae and maxillary palps (Figure S3). The remaining responses likely originate in the gustatory organs as gustatory receptor neurons also directly respond to odors (Wei et al., 2022). Odor responses increased with starvation and plateaued after 4 h (Figures S4), which coincides with the time when output from olfactory receptor neurons (ORNs) plateaus (Root et al., 2011). In sum, DANs exhibit stereotypical responses to a variety of odors.

### DANs encode the innate value of odors

Examination of responses to representative attractive and aversive odors indicated that DANs in individual compartments encode the innate value of odors. For example, α’2, α’1, γ1, γ2, and γ3 compartments preferentially responded to aversive odors whereas γ4, β1, and β’2 compartments preferentially responded to attractive odors (Figure 2A). Several compartments also showed odor value-dependent differences in response dynamics. α3 and α’2 compartments responded more strongly to aversive as compared to attractive odors at stimulus onset but this relationship was flipped at stimulus offset (Figure 2B), suggesting that these DANs signal the removal of attractive stimuli as an aversive event. The opposite dynamics were observed in β’2 compartment (Figure 2B).

**Figure 2.**
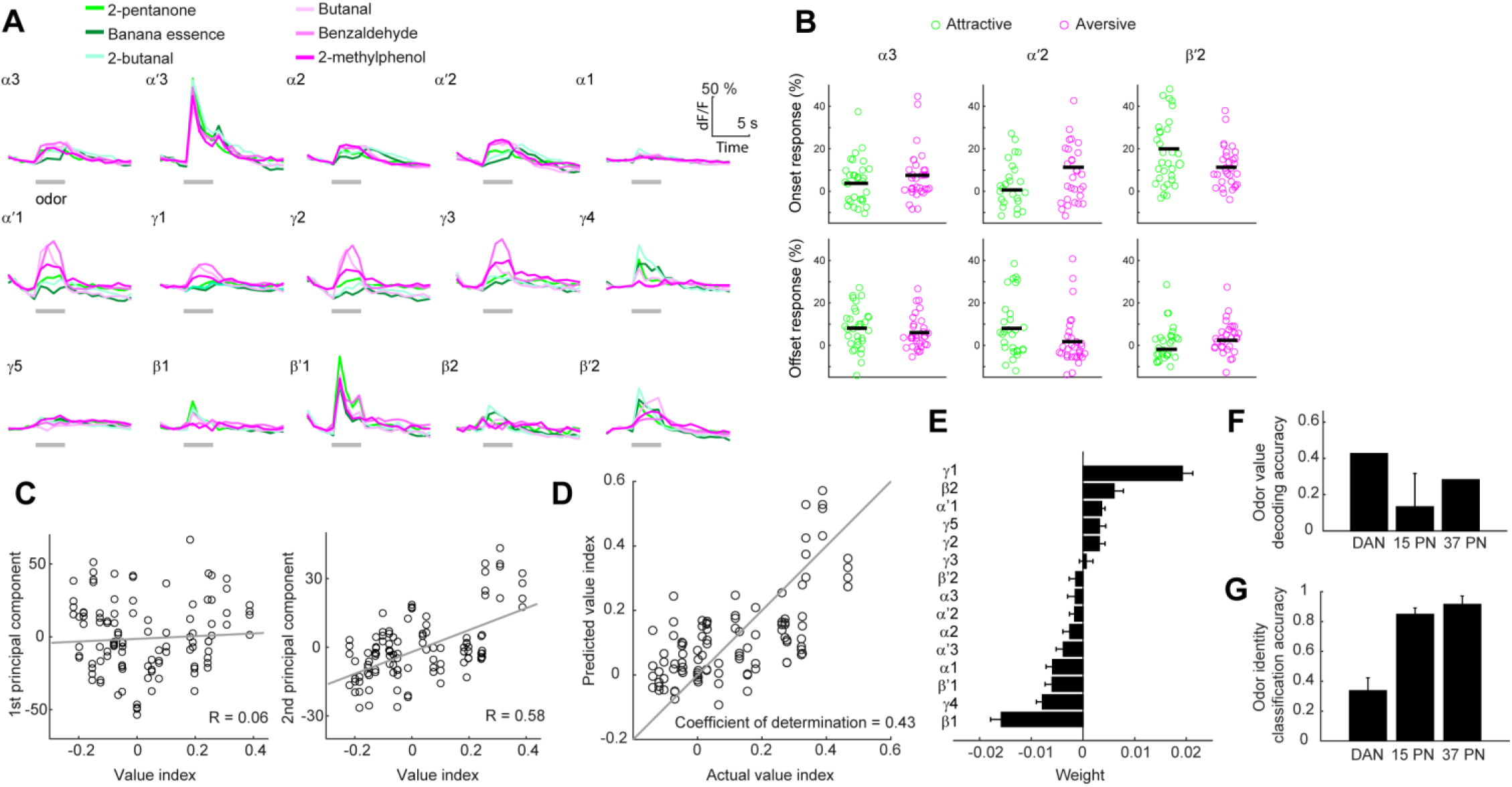
DANs encode innate value of odors (A) Average DAN responses to the three most attractive and aversive odors (average across trials and flies). Gray bar indicates the period of odor application (4 s). (B) Onset responses (average across 2 time frames following odor onset and across trials) and offset responses (average across 2 time frames following odor offset and across trials) to the three most attractive (green) and aversive (magenta) odors. Each dot represents an odor-fly pair. (C) Characterization of DAN population responses to the 27 odors with principal component analysis (average across 3 time frames and across flies). Each dot represents an odor-trial pair. The 2^nd^ principal component is correlated with the value index (R = 0.58, p = 7.3×10^−11^, variance explained = 59 and 19 % for 1^st^ and 2^nd^ principal component). (D) Linear models predict odor values from DAN responses (Coefficient of determination = 0.43). The value of odor not used in training was predicted for each model. Each dot represents an odor-trial pair. Data comprises responses to all the odors except for the controls (water and air). (E) Weights for individual compartments in the final decoding model (an average across 24 sets of regression weights obtained in cross-validation). Negative and positive weights correspond to contribution to attractive and aversive odor values, respectively. (F) Accuracy of odor value prediction made with responses of DANs and PNs. Accuracy of prediction was quantified with coefficient of determination as in D. To compare the performance with the same number of explanatory variables, 15 out of 37 PN types were randomly chosen (n = 100 trials). The odor value was more accurately predicted with responses of DANs as compared to those of PNs (p = 5.2×10^−29^ for DAN vs 15 PN types, one-sample t-test). (G) Accuracy of odor identity prediction made with responses of DANs and PNs. Data were randomly divided into 4 groups for cross-validation, which was repeated for 100 times. Odor identity was more accurately predicted with responses of PNs as compared to those of DANs (p = 9.1×10^−123^ for DAN vs 15 PN types, two-sample t-test). Error bar represents SD. DAN data are from the same 10–11 flies as in Figure 1 and PN data are from Badel et al. (2016).

To understand the coding at a population level, we applied principal component analysis to odor responses of 15 compartments. We found that the second principal component correlates with the odor value, capturing 19% of the total response variance (Figure 2C). We further asked whether odor value can be decoded by a weighted sum of the activity of DANs in 15 compartments. We trained the model using partial least-squares regression and tested if it could predict the value of odor that was not used in training (see Methods). The predictive power of the model was significantly higher than that of controls (Figure 2D, coefficient of determination = 0.43 and − 0.36 ± 0.21 (n = 100 shuffles) for the actual vs odor value shuffled control models, p = 1.5×10^−59^, one sample t-test), demonstrating that DANs encode information about the innate value of odors. The distribution of weights indicates that individual compartments differentially contribute to value coding (Figure 2E).

We previously reported that innate odor values can also be decoded from PNs (Badel et al. 2016). This prompted us to examine whether DANs that are further downstream in the circuit encode odor values more accurately than PNs. The decoding analysis showed that the odor value was indeed more accurately predicted with responses of DANs as compared to those of 15 randomly chosen PN types (to match the number of MB compartments) or even all 37 PN types recorded in the previous study (Figure 2F; Badel et al. 2016). In contrast, the odor identity was more accurately predicted by a model with responses of a subset or all the recorded PNs as compared to those of DANs (Figure 2G). Thus, the olfactory information encoded in PNs is transformed to emphasize the value over identity of odors as it reaches DANs.

### A network model based on the connectome data recapitulate the characteristics of DAN responses

To gain insights into the circuit mechanism underlying PN to DAN olfactory transformation, we turned to the connectome data constructed from electron microscopy images (Clements et al., 2020; Scheffer et al., 2020). We extracted all the pathways between PNs and DANs connected via one or two interneurons and built a neural network comprising neurons within these pathways (Figure 3A, see Methods). Interneurons could be classified into three groups; group 1 receives input from PNs and other interneurons and sends output to other interneurons or DANs; group 2 receives input from PNs and sends output to other interneurons; group 3 receives input from other interneurons and sends output to DANs. Group 1 mainly consists of KCs and forms recurrent connections among group members, whereas group 2 contains lateral horn neurons (LHNs), and group 3 contains MBONs as well as neurons innervating regions including crepine and superior medial protocerebrum (Figure 3B). The network suggests that both the MB and LH pathways convey olfactory information to DANs with the former accommodating a recurrent besides feedforward circuit motif (Figure 3A).

**Figure 3.**
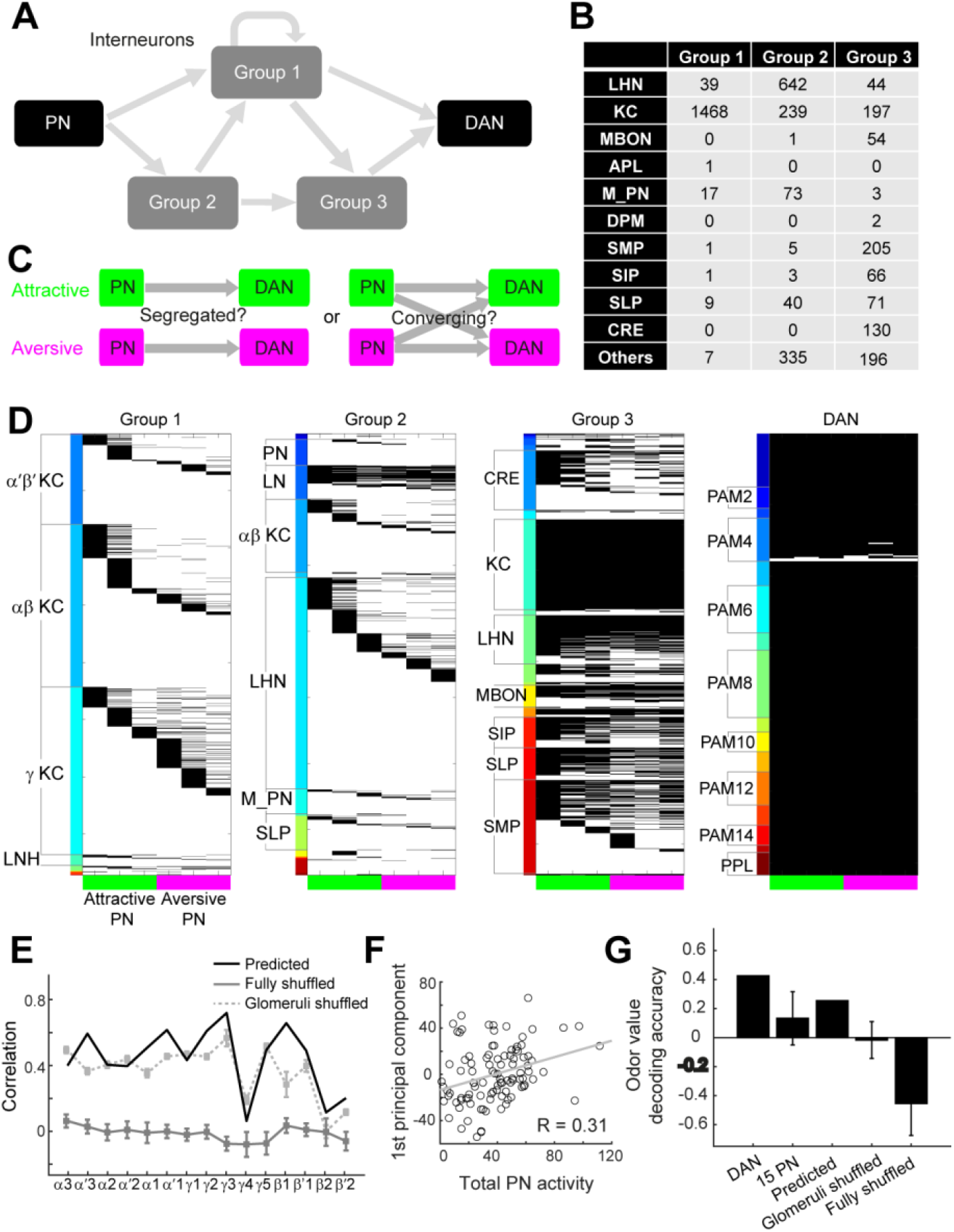
A network model based on the connectome data recapitulate the general aspects of DAN responses (A) Schematic of a network architecture constructed from the connectome data. Interneurons between PNs and DANs could be classified into three groups (see Results). (B) Neuron types in each interneuron Group. Table indicates the number of each type of neurons. Abbreviations: APL (anterior paired lateral), DPM (dorsal paired medial), M_PN (multi-glomerular PNs), SMP (superior medial protocerebrum), SIP (superior intermediate protocerebrum), SLP (superior lateral protocerebrum), and CRE (Crepine). (C) Schematic of two hypotheses on the connectivity between attraction- and aversion-contributing PNs and DANs. (D) Propagation of signals originating in two sets of PNs that contribute most strongly to attraction or aversion. PNs were selected according to Badel et al., 2016, whose identities are DM4, DM1, VA1v, DC3, VA6, and DL5 from the left to right. Interneurons are grouped by their identity and sorted by their connection with PNs. Interneurons that receive signals originating in the corresponding PNs are indicated with a black bar. Most of the DANs reside in two clusters called protocerebral posterior lateral 1 (PPL1) and protocerebral anterior medial (PAM). PPL1 neurons project primarily to the vertical lobes and convey negative reinforcement signals during learning (Aso et al., 2010, 2012; Claridge-Chang et al., 2009; Mao and Davis, 2009). PAM neurons project primarily to the medial lobes and convey positive reinforcement signals (Burke et al., 2012; Liu et al., 2012). (E) Correlation between the actual and predicted DAN responses calculated with the original data (predicted, black), after shuffling the glomerular and odor identities (fully shuffled, dark gray), or after shuffling only the glomerular identity (glomeruli shuffled, light dotted gray). Data comprises responses to all the odors except for the controls (water, mineral oil, and air). Average odor responses (across 4 trials) are used to calculate the correlation. (F) 1^st^ principal component of population DAN responses is correlated with total PN responses to the same set of odors (R = 0.31, p = 0.0018). (G) Accuracy of odor value decoding based on actual DAN responses, actual PN responses (15 glomeruli randomly selected, 15 PN), predicted DAN responses, glomeruli shuffled DAN responses, or fully shuffled DAN responses. Random selection of glomeruli and shuffling were conducted for 10 times. Accuracy was assessed using coefficient of determination as in Figure 2D. Accuracy of decoding with predicted DAN responses is significantly higher than that with PN responses or shuffled DAN responses. (p = 2.3×10^−9^, 1.2×10^−4^, and 4.6×10^−6^, for 15 PN, glomeruli shuffled, and fully shuffled, one-sample t-test with Bonferroni correction). Error bar represents SEM. DAN data are from the same 10–11 flies as in Figure 1 and PN data are from Badel et al. (2016).

We have previously shown that PNs in individual glomeruli contribute to either attraction or aversion to odors (Badel et al., 2016). To start to address whether these attraction- and aversion-contributing PNs form segregated or converging pathways to shape odor value representations in DANs (Figure 3C), we traced the circuit and asked whether each neuron in the network received input from two sets of three PNs that most strongly contribute to attraction or aversion (Figure 3D). Input from attractive and aversive PNs were mostly segregated in groups 1 and 2, with a larger number of α’/β’ KCs, α/β KCs, and LHNs preferentially receiving input from attractive PNs, and γ KCs preferentially receiving input from aversive PNs. However, these PN pathways gradually converged such that the majority of group 3 neurons and almost all the DANs received input from both types of PNs (Figure 3D), suggesting that DAN activity is generated through integration of PN inputs through dense, yet specific connectivity with proper weights.

We therefore asked whether odor responses of DANs can be predicted from those of PNs (Badel et al., 2016) in this network where synaptic connectivity and weights, reflecting the number of synapses, are fully constrained by the connectome (Scheffer et al., 2020). The sign of the output was set to excitatory or inhibitory according to the accumulated evidence provided in the open database (Clements et al., 2020). Neuronal output was determined by a weighted sum of input followed by nonlinear rectification. Using the synaptic weights in the database (i.e. without further tuning the synaptic weights), our model predicted responses that are similar to those observed in our experiments (R = 0.45, the correlation coefficient between the actual and predicted responses over all the compartments; R = −0.027 ± 0.08, control with shuffled glomerular identity; p = 2.2×10^−08^, one-sample t-test). The correlation was significantly higher than chance also for individual compartments albeit with some variability (Figure 3E). Although the difference was less pronounced as compared to another control, where only the glomerular identity was shuffled (glomeruli shuffled), this was to be expected as activity of PNs in different glomeruli are correlated to some extent (Bhandawat et al., 2007), and DANs encode total PN activity across glomeruli besides odor values (Figure 3F).

Critically, odor values could be decoded from predicted DAN responses with accuracy higher than from PN responses or fully shuffled DAN responses (Figure 3G). The decoding accuracy was also higher than the case where only the glomerular identity was shuffled (Figure 3G). Our simulation therefore suggests that information about odor values is extracted, in part, through specific connectivity embedded in a network bridging PNs and DANs.

### DANs integrate multisensory values

Having characterized the representation of odor value in DANs, we next sought to investigate how this relates to the representation of well-described gustatory value and how values associated with different modalities are integrated in DANs. To this end, we compared the responses of DANs to olfactory and gustatory stimuli applied either individually or simultaneously. Because it is technically challenging to apply a tastant with high temporal precision, and an intake of a tastant could cause satiation leading to a change the internal state of an animal, we attempted to replace tastant application with optogenetic activation of sweet- or bitter-sensing gustatory receptor neurons (GRNs; Figures 4A and S5). Activation of Chrimson-expressing Gr64f and Gr66a GRNs produced a response resembling that to sucrose and quinine, respectively (Figure 4B, C; Pearson’s R is 0.78 for Gr64f GRN stimulation vs sucrose, and 0.84 for Gr66a GRN stimulation vs quinine). Using this technique, we applied olfactory and gustatory stimuli individually and found that responses to an aversive odor (2-methylphenol) and bitter GRN activation were well correlated across compartments (Figure 4D; Pearson’s R = 0.84, p = 7.7×10^−05^). On the other hand, responses to an attractive odor (2-pentanone) and sweet GRN activation were less correlated, with some compartments (γ5 and α’3) mainly responding to one of the two stimuli (Figure 4D; Pearson’s R = 0.02, p = 0.95). Thus, negative values are represented similarly regardless of sensory modalities whereas positive values bear some modality specificity.

**Figure 4.**
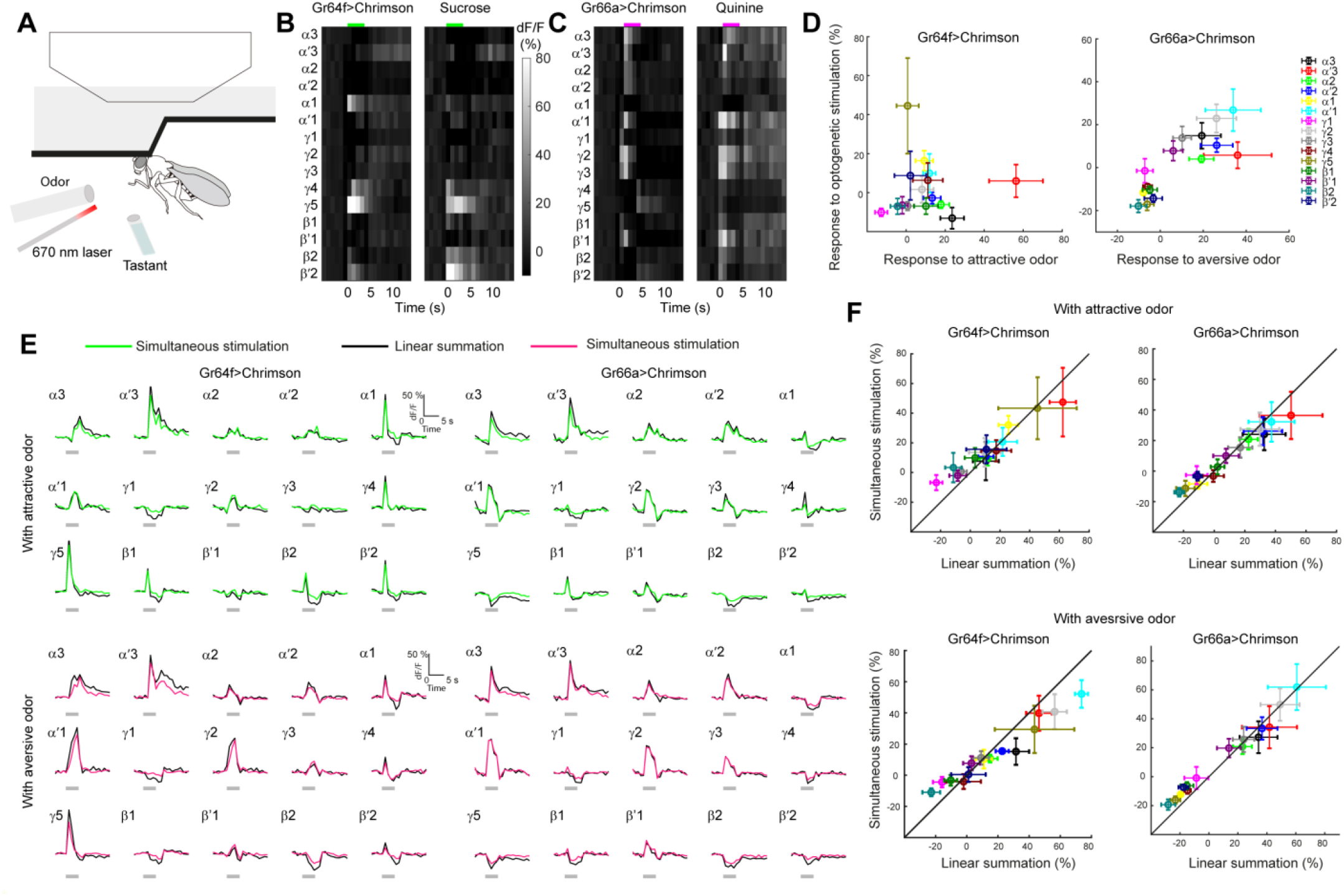
DANs integrate multisensory values (A) Setup for imaging DAN responses to odors, tastants, or optogenetic stimulation of GRNs in isolation or in combination. (B) Responses of DANs to optogenetic stimulation of Chrimson-expressing Gr64f GRNs or sucrose (average across trials and flies, n = 6 flies). Pearson’s correlation between the two responses is 0.78 (p = 5.6×10^−4^). Colored bar at the top indicates the period of stimulation (4 s). (C) Same as in B but for optogenetic stimulation of Chrimson-expressing Gr66a GRNs or quinine (average across trials and flies, n = 5 flies). Pearson’s correlation between the two responses is 0.84 (p = 7.4×10^−5^). (D) Comparison between gustatory (optogenetic) and olfactory responses of DANs (averaged across time and trials, n = 6 flies) in individual compartments. (E and F) Integration of gustatory (optogenetic) and olfactory responses by DANs. (E) Comparison between the sum of DAN responses to individual stimulation (black) and DAN responses to simultaneous application of gustatory and olfactory stimuli (green, attractive odor; magenta, aversive odor; average across trials and flies, n = 6 flies). Attractive and aversive odors were paired with both Gr64f and Gr66a GRN stimulation. Gray bar indicates the period of stimulus application (4 s). (F) The same data as in E, but for comparison between peak responses in individual compartments. The same color code as in D. Error bar represents SD.

We next applied olfactory and gustatory stimuli simultaneously and compared the response to the linear sum of individual responses (Figure 4E). We paired both attractive and aversive odors with a sweet or bitter stimulus. We found a good match between simultaneous stimulation and linear summation of individual stimulation in terms of amplitude and kinetics (Figure 4E). This trend was observed across 15 compartments (Figure 4F), demonstrating that DANs respect innate olfactory values just as much as gustatory values during multimodal integration.

### Repetitive application of odors induces compartment- and value-dependent plasticity in MBONs

The observation that odors recruit DANs concomitantly with sensory neurons raises the possibility that odors alone induce value-dependent plasticity in the MB. Previous studies have shown that, within a compartment, DANs responding to attractive stimuli such as sucrose are coupled with MBONs that drive avoidance behavior (Aso et al., 2014b; Burke et al., 2012; Liu et al., 2012). DANs induce depression at KC to MBON synapses when they are activated together with KCs (Handler et al., 2019; Hige et al., 2015; Owald et al., 2015), resulting in weaker transmission of an avoidance signal from this MBON. Therefore, sampling of an attractive odor is expected to increase attractive drive to that odor (Figure 5A). Because compartments innervated by DANs responding to aversive stimuli such as an electric shock are coupled with MBONs that drive attraction (Aso et al., 2010, 2014b; Claridge-Chang et al., 2009; Schwaerzel et al., 2003) a mirror symmetric phenomenon is expected to take place upon sampling of an aversive odor (Figure 5E). Thus, we hypothesized that odors alone can induce plasticity in MBONs depending jointly on the value of odor and the identity of compartment.

**Figure 5.**
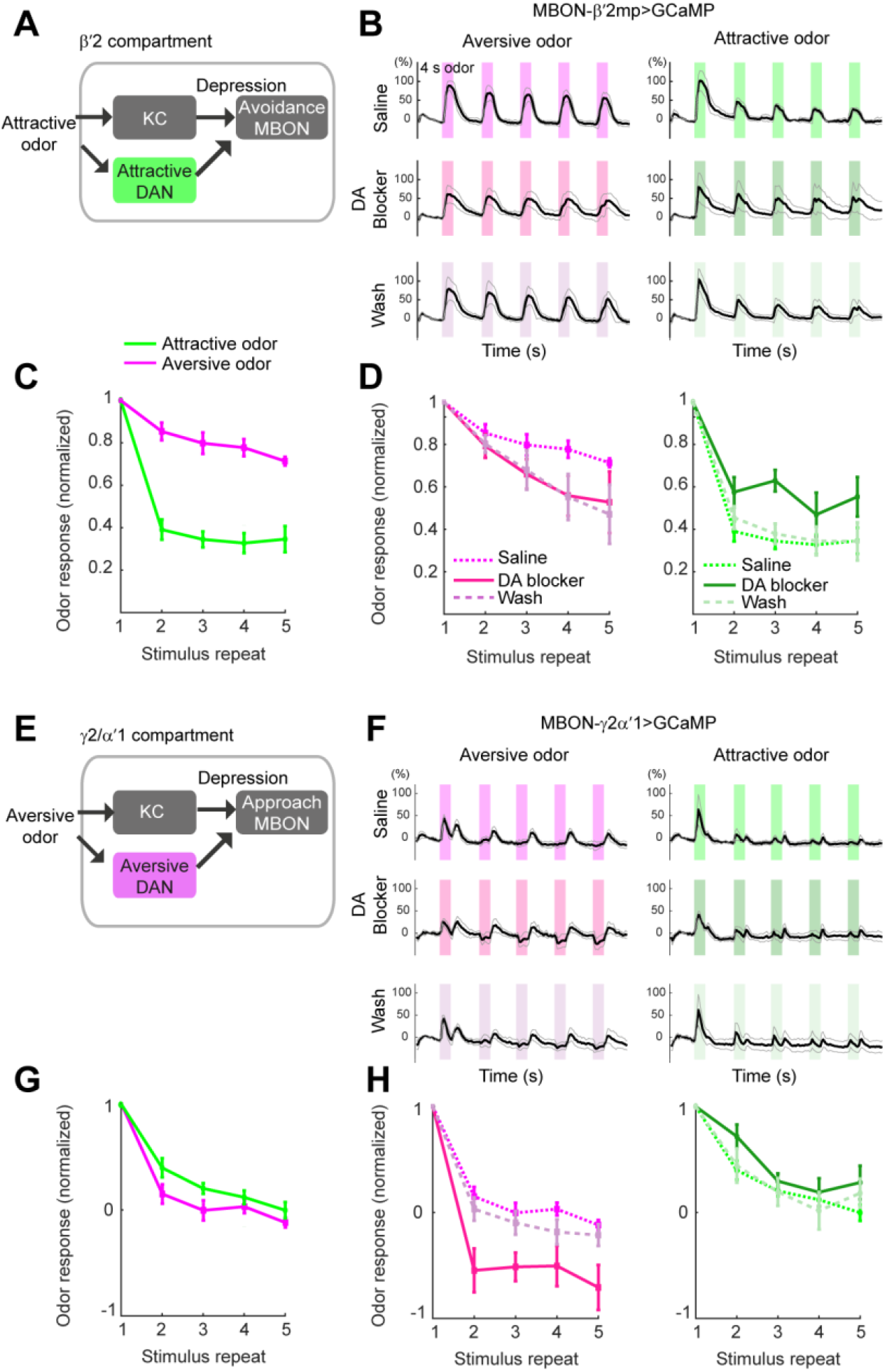
Repetitive application of odors induces compartment- and value-dependent plasticity in MBONs (A) Schematic of a working model in β’2 compartment, where attractive odors activate DANs preferentially besides KCs and induce depression in avoidance-driving MBON-β’2mp activity. (B) Responses of MBON-β’2mp to repetitive application of attractive (2-pentanone) or aversive (2-methylphenol) odors in saline, after the addition of a blocker of dopamine receptor (10 μM SCH 23390), or after wash (average across trials, n = 5 flies). Colored region indicates the period of odor application (4 s). Black trace represents the mean and gray traces represent the range of SEM. (C) Peak responses (average across 8 time frames after odor onset and across trials) of MBON-β’2mp to attractive and aversive odors in saline, normalized to the response to the first stimulus. Responses are more depressed in response to an attractive over aversive odor (comparison between responses to two different odors in saline, p = 1.5×10^−6^, two-way ANOVA). (D) Normalized responses of MBON-β’2mp in saline, SCH 23390, or after wash. SCH 23390 more strongly modulated the responses to an attractive over aversive odor (p = 5.1×10^−7^ and 0.066 for attractive and aversive odors, respectively, two-way ANOVA). (E-H) Same as in A-D, but for responses of MBON-γ2α’1 in γ2/α’1 compartments (n = 6 flies). (F, G) Responses are more depressed in response to an aversive over attractive odor (comparison between responses to two different odors in saline, p = 0.0028, two-way ANOVA). (H) SCH 23390 more strongly modulated the responses to an aversive over attractive odor (p = 1.1×10^−7^ and 0.057 for aversive and attractive odors, respectively, two-way ANOVA). Because the responses were more transient in this compartment, peak responses were calculated as an average across 4 time frames after odor onset and across trials. Error bar represents SEM. P values for all statistical tests are listed in Table S4.

To test this, we recorded the responses of MBONs in β’2 and γ2/α’1 compartments (MBON-β’2mp and MBON-γ2α’1, respectively; Aso et al., 2014b) to repetitive application of attractive and aversive odors (Figure 5B, F). In β’2 compartment, because DANs are more strongly activated by attractive odors (Figure 2), stronger depression likely occurs in MBONs in response to attractive over aversive odor application. Indeed, responses to an attractive odor depressed more rapidly (Figure 5B, C). The same trend was obtained with a different pair of attractive and aversive odors, further indicating that the differential plasticity is odor value-rather than identity-dependent (Figure S6). In a similar vein, in γ2/α’1 compartments where DANs are activated more strongly by aversive odors (Figure 2), stronger depression occurred in response to an aversive odor (Figure 5F, G), demonstrating that odor value-dependent plasticity takes place in multiple compartments.

To investigate if this plasticity requires the action of dopamine, we examined the effect of SCH 23390, a D1 type dopamine receptor antagonist (Boto et al., 2014), on MBON responses. We noticed that the antagonist reduced the overall magnitude of responses, but it also affected the plasticity induced by repetitive application of odors (Figure 5B, D, F, H). In β’2 compartment, the antagonist reversibly reduced the extent of MBON depression evoked by an attractive but not an aversive odor (Figure 5D), consistent with attractive DANs being a partner in this compartment. In γ2/α’1 compartments, the antagonist modulated the plasticity evoked only by an aversive odor, again, matching the property of the cognate DANs (Figure 5H), although the effect was more complex including the emergence of an inhibitory response (Figure 5F), implying that dopamine has multiple sites of action in shaping the response of MBONs in this compartment. These results indicate that odor-induced plasticity in MBONs is mediated by dopamine.

### DANs drive odor value-dependent modulation of navigation

Previous studies have shown that MBONs in each compartment promote either approach or avoidance and the total activity of MBONs across compartments determines the final behavioral output (Aso et al., 2014b; Owald et al., 2015). According to this model developed in the context of associative learning, modulation of MBON activity by DANs biases the behavior to the side determined by the identity of MBONs (Aso et al., 2014b; Owald et al., 2015). Here we have shown that DANs acutely modulate MBON activity in the context of olfactory sensory experience (Figure 5), suggesting that DANs also shape the behavior of the fly as it navigates an olfactory environment. To test this hypothesis, we let the flies sample odors in a flight simulator (Figure 6A; Badel et al., 2016) and examined if their responses to odors were affected by inhibition of specific types of DANs through optogenetic activation of an anion channel, GtACR1 (Govorunova et al., 2015; Mohammad et al., 2017). The flight simulator was run in closed-loop, such that the tethered fly was able to control the visual and olfactory environment by adjusting its virtual heading direction through turns detected by the difference in the left and right wingbeat amplitudes (Figure 6A; Badel et al., 2016). Odors were provided in the form of a plume spanning 45 degrees in azimuth. We used three odors with different values (neutral, aversive, attractive), and each odor was provided three times in succession with fixed intervals before switching to the next odor (Figure 6B). Flies rapidly escaped from an aversive odor by making sharp turns whereas they flew straight and spent more time inside an attractive odor (Figure 6D). We quantified the behavioral responses to an odor by calculating the proportion of time the fly has spent outside of an odor (Figure 6E). DANs were optogenetically inhibited only when the fly was within the odor plume (Figure 6B).

**Figure 6.**
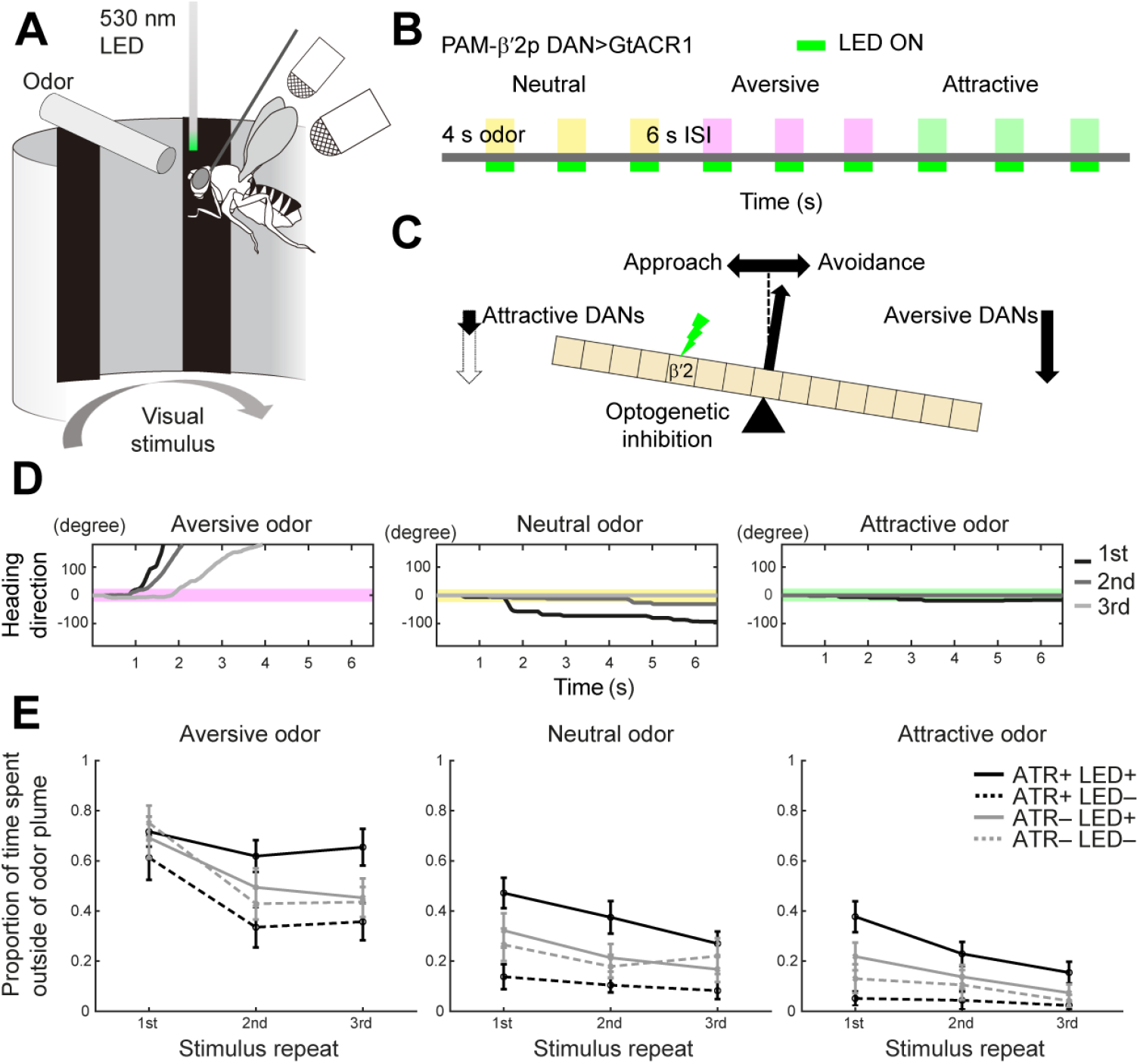
DANs drive odor value-dependent modulation of navigation (A) Schematic of the virtual-flight arena. (B) Schematic of the behavioral assay. Three odors with different values (γ-butyrolactone, neutral; 2-pentanone, attractive; 2-methylphenol, aversive) were each applied three times in succession with a 6 s inter-stimulus interval. 530 nm LED light was applied when the fly was in the odor region spanning 45 ° in azimuth. The sequence of stimulation was repeated for 7 times. Because the experiment was run in closed-loop, flies had a choice of staying in or outside of the odor region during the 4 s stimulus application period. (C) Schematic of a working model explaining the effect of suppressing DAN activity on the olfactory behavior of flies. In control condition, the effects of attractive DANs coupled with avoidance-driving MBONs and aversive DANs coupled with approach-driving MBONs are balanced, but the suppression of attractive DANs will relax odor-evoked depression in avoidance-driving MBONs and tip the behavior to aversion. (D) Trajectories of a representative fly in response to three repeated applications of three odors. Colored areas indicate the odor region (45 °). (E) Proportion of time that flies spent outside of an odor for each stimulus repeat. The experiment was conducted under four different conditions, in which ATR and LED were applied (+) or not (−) (ATR+LED+, ATR+LED −, ATR − LED+, ATR − LED −; n = 20–23 flies for each condition). Optogenetic suppression of PAM-β’2p DANs significantly biased the behavior towards aversion in response to all three odors (p = 0.0039, 1.1×10^−6^, and 1.2×10^−6^, for aversive, neutral, and attractive odors, two-way ANOVA). Error bar indicates SEM. P values for all statistical tests are listed in Table S4.

We first targeted PAM-β’2p DANs, a partner of MBON-β’2mp that are shown to drive avoidance behavior (Figure 5A; Aso et al., 2014a, 2014b). Our physiological experiments (Figure 5) suggest that suppression of PAM-β’2p relaxes odor-evoked depression in MBON-β’2mp, thus biasing the behavior towards avoidance (Figure 6C). As expected, the flies avoided the odors more strongly under the presence of light and all-trans-retinal (ATR), an essential cofactor of GtACR, as compared to controls (Figure 6E). In particular, responses to an aversive odor diverged more clearly over repeated stimulation; the level of avoidance remained high under the presence of light and ATR. This is likely because suppression of DAN-induced plasticity counteracts the effect of adaptation. Little behavioral effect was observed following the suppression of PPL1-γ2α’1 DANs (Figure S7), presumably reflecting a complex effect of dopamine on MBONs in this compartment (Figure 5F). The data in β’2 compartment, however, reveals a causal role of DANs in moment-to-moment control of olfactory navigation.

## Discussion

What makes DANs unique is their versatile functions exerted under different behavioral contexts, a feature common across species (Cohen et al., 2012; Engelhard et al., 2019; Howe et al., 2013; Schultz et al., 1997; da Silva et al., 2018). In *Drosophila*, a large body of work has shown that DANs drive learning (Aso and Rubin, 2016; Aso et al., 2010, 2019; Burke et al., 2012; Claridge-Chang et al., 2009; Handler et al., 2019; Hattori et al., 2017; Hige et al., 2015; Kim et al., 2007; Liu et al., 2012; Owald et al., 2015; Plaçais et al., 2012; Schwaerzel et al., 2003; Yamagata et al., 2015), mediate memory extinction, reconsolidation, and forgetting (Berry et al., 2012, 2015; Felsenberg et al., 2017, 2018; Shuai et al., 2015) exert internal state-dependent behavioral control (Alekseyenko et al., 2013; Kaun et al., 2011; Krashes et al., 2009; Sayin et al., 2019; Tsao et al., 2018; Ueno et al., 2012; Yamagata et al., 2016) and represent movement (Cohn et al., 2015; Siju et al., 2020; Zolin et al., 2021). Here we introduce an additional function of DANs: dynamic updating of sensory values during navigation.

### Encoding and integration of multisensory values by DANs

In associative conditioning, DANs are regarded to encode information about the reinforcer, whose value is conferred to a conditioned stimulus encoded by separate sensory neurons (Schultz et al., 1997; Spanagel and Weiss, 1999; Watabe-Uchida et al., 2017). However, our results show that this view of mutually exclusive encoding needs to be re-considered because DANs also respond to odors that are commonly used as a conditioned stimulus. Specifically, DANs in individual compartments differentially responded to odors with opposing values and the decoding analysis demonstrated that a population of DANs carry information about the innate value of odors (Figure 2). A recent study using 7 odors that are mostly non-overlapping with our odor set showed that DANs in a similar set of compartments as we describe here encode odor valence (Siju et al., 2020), suggesting the generality of this coding property. Moreover, we found that DANs encode odor values more accurately than PNs. These lines of evidence suggest that the extent of olfactory conditioning is jointly determined by the value of reinforcer and the innate value of co-applied odors that together set the activity of DANs. It also indicates that odors with large innate values themselves may function as reinforcers when they coincide with visual or gustatory signals entering the MB (Kirkhart and Scott, 2015; Vogt et al., 2014, 2016; Yagi et al., 2016).

How do these representations of odor values in DANs compare with those of better characterized taste values? We found a strong correlation between responses to an aversive odor and bitter GRN activation, whereas weaker correlation was discerned between responses to an attractive odor and sweet GRN activation. The former relationship likely reflects the ethological importance of unequivocally recognizing any cues that are predictive of negative consequences. On the other hand, the latter indicates that DANs appreciate different aspects of positive values inherent in different sensory modalities. Therefore, cues with positive values are expected to induce plasticity in specific sets of MB compartments depending on their sensory modality.

Although studies have reported DAN’s responses to salient cues of different modalities (Kirkhart and Scott, 2015; Liu et al., 2012; Riemensperger et al., 2005; Tomchik, 2013), little attempts have been made to characterize how they are integrated. This issue is particularly relevant under natural settings as sensory experiences such as feeding arguably involve multimodal inputs (Spence, 2015). Our results showed that gustatory and olfactory values are integrated linearly in DANs across compartments. This was true for all four combinations of positive and negative stimuli of two modalities, illustrating that DANs arithmetically treat values of different modalities and signs equally, and serve as a major node of value integration.

### Structure of circuits underlying odor value coding by DANs

To decipher the circuit mechanisms underlying multifaceted functions of DANs, previous studies in rodents and flies have identified neurons providing direct input to DANs (Beier et al., 2015; Lerner et al., 2015; Menegas et al., 2015; Otto et al., 2020; Watabe-Uchida et al., 2012). Neural activity in these presynaptic neurons has also been systematically measured to examine how DANs compute reward prediction errors (Tian et al., 2016). However, characterization of multilayered circuits connecting all the way from sensory neurons to DANs has remained a challenge. Here we utilized the fly connectome data generated based on electron microscopy (Scheffer et al., 2020) and analyzed the circuit bridging PNs and DANs. We found that both the MB and the LH pathways converged on DANs with the former harboring feedback as well as feedforward motifs. In terms of odor value processing, PNs implicated most strongly in attraction and aversion (Badel et al., 2016; Semmelhack and Wang, 2009) differentially innervated certain interneuron types in earlier layers. However, these inputs increasingly converged in deeper layers and DANs, suggesting that odor value is computed in DANs by integrating the activity originating from diverse PN types.

A wiring diagram reflecting experimentally determined synaptic connections is a major piece of information required for understanding the mechanism of neural processing, and surging attempts are made to mine the statistical properties of connections (Li et al., 2020; Otto et al., 2020; Scheffer et al., 2020; Schlegel et al., 2021; Zheng et al., 2018). However, it is still one of the many necessary pieces, and it has not been examined to what extent the connectome can recapitulate the activity of output neurons from that of distant, input neurons when combined with core biological properties. To address this issue, we have capitalized on our network model and comprehensive data set of odor responses in both the input and the target layers, namely, PNs (Badel et al., 2016) and DANs (Figure 1). Because PN responses can be predicted from the responses of olfactory receptor neurons (Olsen et al., 2010), our analysis effectively connects all the way from the olfactory input to DANs. We incorporated synaptic integration with reported sign of synapses (Clements et al., 2020) as well as neuronal nonlinearity reflecting spiking into the network model, and found that the model recapitulated aspects of DAN responses including the coding of odor values and odor-specific total PN activity. A larger amount of information about odor values was encoded in simulated DANs as compared to PNs.

As the extent of recapitulation was limited, there is much room for improvements such as considering the pathways connecting PNs and DANs with more than three interneurons, determining the sign of synapses for larger number of neurons, characterizing the short-term plasticity of individual synapses, and probing intrinsic properties of individual neuronal types. Nevertheless, our simulation with physiological input and output provides a way to identify the circuit motifs that are pivotal for actual functions.

### Dynamic modulation of neuronal output and olfactory navigation by DANs

Coding of odor values by DANs has consequences beyond the context of associative conditioning. Because odors alone recruit sensory and DAN pathways simultaneously, DAN-dependent plasticity is expected to occur in the MB compartments and consequently modulate the behavior while the fly navigates the olfactory environment. Importantly, this plasticity should depend on both the identity of DANs and the value of odors. Indeed, we found that in β’2 compartment innervated by attractive DANs, an attractive odor induced much stronger depression in MBON-β’2mp responses as compared to an aversive odor. A mirror symmetric phenomenon was observed in γ2/α’1 compartments innervated by aversive DANs. These changes were dependent on dopamine as the blocker of D1 type dopamine receptor antagonist altered the plasticity only in response to an odor that preferentially activated DANs in respective compartments. Because the blocker revealed an additional inhibitory input to MBON-γ2α’1, there likely exist yet uncharacterized interactions involving DANs and inhibitory MBONs innervating these compartments such as MBON-γ4>γ1γ2 (Aso et al., 2014a).

Consistent with the effects on MBONs, odors evoked DAN-dependent modulation in behavior as the fly navigated the virtual olfactory environment. Specifically, optogenetic suppression of attractive PAM-β’2p DANs made the responses to odors more aversive, demonstrating a causal role of DANs in the moment-to-moment control of value-dependent sensory exploration. This behavioral modulation is adaptive as sampling of an attractive odor will tend to further enhance the drive toward this odor through the activation of attractive DANs and subsequent suppression of avoidance MBONs.

Recent studies support this modulatory role of odors in other behavioral contexts. For example, olfactory experiences form a short-lived memory that alter the performance of subsequent olfactory association, an effect named latent inhibition (Jacob et al., 2021). This requires the output of γ2α’1 and α3 DANs in case of inhibiting appetitive memory. In another context of active odor pursuit, γ4 DANs that multiplex information about an odor and self-movement help walking flies track apple cider vinegar (Zolin et al., 2021). Because γ2α’1 and α3 DANs are of aversive and γ4 DANs are of attractive type, the direction of behavioral biases observed in these studies conforms with our finding of odor value-dependent regulation by DANs. In mice, responses of DANs to novel but not familiar odors are correlated with the innate attractiveness of odors and promote subsequent associative learning (Morrens et al., 2020), bearing similarity to the effect in flies. These and our results therefore point that DANs are in action even in between reinforcing events. As animals in a natural environment sample a barrage of sensory cues with innate values and novelty, DANs continuously update the value of currently experiencing sensory cues and also affect the way sensory values are re-evaluated in a future association. This provides a view that DANs take into account the history of not only canonical reinforcers but also all the continuously experienced external input with values to shape the meaning of sensory cues.

## Acknowledgments

We thank Adam Claridge-Chang, Michael Dickinson, André Fiala, Vivek Jayaraman, Akinao Nose, Gerald Rubin, Hiromu Tanimoto, and Bloomington Stock Center for fly stocks; Yoshinori Aso for the template 3D data of the MB compartments; Henrik Skibbe for advice on non-linear image registration; Takuya Isomura for comments on the connectome analysis. We are grateful to Naoshige Uchida and Kenji Morita for comments on the manuscript, and members of the Kazama laboratory for their support and comments on the manuscript. This work was funded by a grant from RIKEN, a grant from Kao Corporation, and JSPS KAKENHI Grants (JP18H02532, JP18K19502, JP21H04789) to H.K. A.K. was supported by RIKEN JRA fellowship, Grant-in-Aid for JSPS Fellows (JP19J12156), and Masason Foundation.

## Author Contributions

A.K. and H.K. designed the study and wrote the manuscript. A.K. performed all the experiments and data analyses except for part of the behavioral experiment, which was performed by K.Ohta.

H.K. and K.Okanoya supervised the project.

## Declaration of Interests

The authors declare no competing interests.

### Methods Fly stocks

Flies (*Drosophila melanogaster*) were raised on a standard cornmeal agar under a 12-h/12-h light/dark cycle at 25°C. All experiments were performed on female flies 3-7 days after eclosion. Flies were starved for 4–7 h with water prior to experiments unless otherwise described. Fly stocks used in this study are as follows: *TH-Gal4* (Friggi-Grelin et al., 2003), *Ddc-Gal4* (Li et al., 2000), *MB247-DsRed* (Riemensperger et al., 2005; gift from André Fiala), *UAS-GCaMP6f(attP40)* (Chen et al., 2013; Bloomington # 42747), *TH-LexA* (Galili et al., 2011; gift from Hiromu Tamimoto), *R58E02-LexA(attP40)* (Liu et al., 2012; Bloomington # 52740), *LexAop-GCaMP6s (attP1)* (Chen et al., 2013; Pfeiffer et al., 2010; Bloomington # 53747), *Gr66a-GAL4* (Kwon et al., 2011; Bloomington #5767043), *Gr64f-Gal4* (Kwon et al., 2011; Bloomington #57669), *UAS-opGCaMP6f, UAS-Chrimson88-tdT(attP18)* (gift from Gerry Rubin), *MB002B-Gal4* (Aso et al., 2014a; Bloomington # 68305), *MB051B-Gal4* (Aso et al., 2014a; Bloomington # 68275), *UAS-CD4::GCaMP6f(attP40); UAS-CD4::GCaMP6f(VK00005)* (Kohsaka et al., 2019; gift from Akinao Nose), *MB056B-Gal4* (Aso et al., 2014a; Bloomington # 68276), *UAS-GtACR1-EYFP(attP2)* (Mohammad et al., 2017; gift from Adam Claridge-Chang), *MB296B-Gal4* (Aso et al., 2014a; Bloomington # 68308). *UAS-GCaMP6f, MB247-DsRed, TH-Gal4, Ddc-Gal4, Gr66a-Gal4, and UAS-GtACR1-EYFP* were backcrossed for six generations to the Dickinson wild type (Dickinson et al., 1999). Detailed genotypes of flies used in each experiment are listed in Table S1.

**Table S1.**
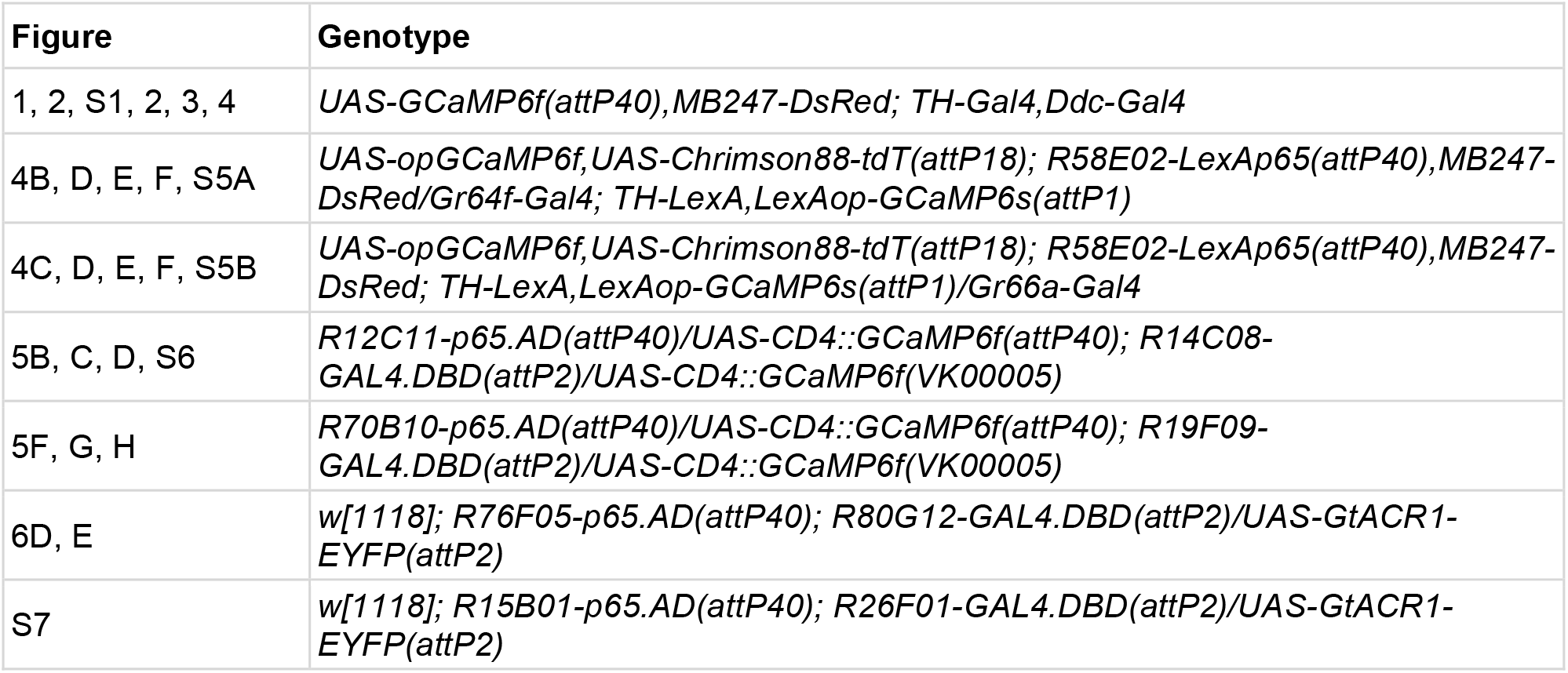
Genotypes used to generate data presented in each figure.

### Olfactory stimulation

Odors were delivered with a custom-made, multi-channel olfactometer controlled by custom MATLAB (MathWorks) code through a data acquisition interface (NI cDAQ-9178, NI 9264 National Instruments) as previously described (Badel et al., 2016). Briefly, an air stream (250 ml/min) was passed through 4 ml of odor solution diluted with mineral oil (nacalai tesque, 23334-85) or water at 1:100 concentration (v/v) except for apple cider vinegar, which was undiluted. The odorized air was further diluted by mixing it with a main air stream (1550 ml/min), a small portion of which was delivered frontally to the fly through a ø2 mm outlet placed 10 mm away from the fly. The speed of odorized air flow at the position of the fly was 0.3 m/s. The detail of odors used in the study is listed in Table S2. The innate odor values are from Badel et al. (2016).

**Table S2.** List of odors used in this study

| Rank | Odor | Innate odor value | Solvent | Vendor |
| --- | --- | --- | --- | --- |
| 1 | 2-methylphenol | 0.466 | oil | Sigma-Aldrich |
| 2 | Benzaldehyde | 0.388 | oil | Sigma-Aldrich |
| 3 | Butanal | 0.336 | oil | nacalai tesque |
| 4 | Acetophenone | 0.325 | oil | Sigma-Aldrich |
| 5 | 1-octanol | 0.322 | oil | Sigma-Aldrich |
| 6 | Hexyl Acetate | 0.277 | oil | Wako |
| 7 | 3-octanol | 0.271 | oil | Tokyo chemical industry |
| 8 | 4-methylcyclohexanol | 0.261 | oil | Sigma-Aldrich |
| 9 | 3-methylthio-1-propanol | 0.18 | oil | Sigma-Aldrich |
| 10 | Methanoic acid | 0.154 | water | Sigma-Aldrich |
| 11 | γ-butyrolactone | 0.13 | oil | Sigma-Aldrich |
| 12 | Linalool | 0.117 | oil | nacalai tesque |
| 13 | Air | 0.079 | NA |  |
| 14 | Ethyl butyrate | 0.065 | oil | Sigma-Aldrich |
| 15 | 1-butanol | 0.029 | water | nacalai tesque |
| 16 | Benzyl Alcohol | 0.028 | water | Wako |
| 17 | Isopentyl Acetate | 0.014 | oil | Wako |
| 18 | Ethyl acetate | 0.002 | oil | Sigma-Aldrich |
| 19 | Mineral Oil | 0 | NA | nacalai tesque |
| 20 | Water | -0.022 | NA |  |
| 21 | Methyl Salicilate | -0.046 | oil | Sigma-Aldrich |
| 22 | Propionic acid | -0.048 | water | Wako |
| 23 | Apple cider vinegar | -0.071 | NA | Mizkan |
| 24 | Pentyl Acetate | -0.074 | oil | Sigma-Aldrich |
| 25 | 2-butanone | -0.104 | water | Sigma-Aldrich |
| 26 | Banana Essence | -0.119 | water | Narizuka corporation |
| 27 | 2-pentanone | -0.138 | oil | Wako |

### Gustatory stimulation

To apply 20 mM sucrose (nacalai tesque, 30403-55) and 10 mM quinine (Sigma-Aldrich, Q1125-5G) to GRNs in the labellum during calcium imaging, the end of a silicone tube was placed ∼2 mm away from the labellum. The other end of the silicone tube was attached to a 1 ml syringe whose position was set with an electronic actuator (KR1501A-0025-H1-000D, THK) controlled by signals sent from MATLAB. The tube and the syringe were filled with a solution. By applying a positive followed by a negative pressure to the solution using the actuator, a drop of solution can be transiently presented and drawn from the outlet of the tube.

To optogenetically stimulate GRNs, Chrimson-expressing Gr64f or Gr66a GRNs were illuminated with a 660 nm LED (M660FP1, Thorlabs). An optic fiber (M118LO3, Thorlabs) connected to the LED through a bandpass filter (FB660-10, Thorlabs) was placed ∼10 mm away from the fly, aiming at the labellum. Flies were fed food with 0.2 mM all-trans-retinal (Sigma-Aldrich, R2500) for at least two days before the experiments.

### In vivo two-photon Ca2+ imaging

Ca^2+^ imaging was conducted as described in (Badel et al., 2016) using a two-photon microscope (LSM 710 NLO, Zeiss) equipped with a water immersion objective lens (W Plan-Apochromat, 20x, numerical aperture 1.0), a piezo motor (P-725.2CD PIFOC, PI), and a titanium:sapphire pulsed laser (Chameleon Vision II, Coherent) mode-locked at 930 nm.

A fly was cold-anesthetized and attached to a custom recording plate with ultraviolet-curing adhesive (NOA 63, Norland). The head cuticle was removed to expose the brain in saline containing 103 mM NaCl, 3 mM KCl, 5 mM N-tris (hydroxymethyl) methyl-2-aminoethane-sulfonic acid, 8 mM trehalose, 10 mM glucose, 26 mM NaHCO3, 1 mM NaH2PO4, 1.5 mM CaCl2, and 4 mM MgCl2 (osmolarity adjusted to 270-275 mOsm) bubbled with 95% O2/5% CO2. Saline was perfused at a rate of 2 ml/min during the recording. The proboscis was fixed in a particular position with ultraviolet-curing adhesive without covering the labellum.

For DAN imaging, DsRed and GCaMP6f signals were recorded simultaneously using two GaAsP detectors through 500-550 nm and 565-610 nm bandpass filters, respectively. 104.61 × 104.61 × 99 µm volume was covered by scanning 34 optical slices separated by 3 μm at x-y resolution of 1.66 × 1.66 µm pixel size, resulting in an acquisition rate of 1.0 s/volume. For MBON imaging, GCaMP signals were recorded using a GaAsP detector through 500-550 nm bandpass filter. 82.36 × 82.46 × 76 µm volume containing the entire dendrites was covered by scanning 20 optical slices separated by 4 μm at x-y resolution of 2.66 × 2.66 µm pixel size, resulting in an acquisition rate of 0.55 s/volume.

To examine odor responses in DANs (Figures 1, 2), each of the 16 odors was applied for 4 s with 60 s inter-trial interval in a block. An experiment consisted of 4 blocks and the order of odor presentation was randomized in each block. Odors were turned on after collecting 5 s of baseline activity.

To compare the responses of DANs to tastants and optogenetic stimulation of GRNs (Figure 4B, C), the former and the latter stimuli each lasting 4 s were applied to the same flies. 20 mM sucrose was applied to Gr64f>Chrimson flies and 10 mM quinine was applied to Gr66a>Chrimson flies. Stimuli were turned on after collecting 10 s of baseline activity.

To examine how olfactory and gustatory inputs are integrated in DANs (Figures 4D-F), odors and optogenetic stimulation of GRNs each lasting for 4 s were applied either independently or simultaneously in the following order and combination with 60 s inter-stimulus interval in a block: attractive odor (2-pentanone), GRN stimulation, simultaneous application of the two, aversive odor (2-methylphenol), GRN stimulation, and simultaneous application of the two. Each experiment consisted of eight blocks. The same set of experiments was conducted for stimulation of Gr64f and Gr66a GRNs.

To examine the responses of MBONs to repetitive application of odors (Figure 5), each odor was applied ten times with 15 s inter-stimulus interval in a trial. Trials corresponding to an attractive (2-pentanone) or an aversive (2-methylphenol) odor were alternated with 2 min intertrial interval in a block, and this was repeated for 4 blocks with 2 min inter-block interval. In each fly, experiment was conducted under three different conditions: under the presence of saline, 10 μM SCH 23390 (Tocris Bioscience), a D1 type dopamine receptor antagonist, and after washing out the antagonist. There was a 5 min interval between conditions.

### Behavioral Experiments

Behavioral responses of flying flies to odors were examined in a flight simulator as described in (Badel et al., 2016). Briefly, the simulator consisted of an olfactometer and an LED visual arena (Mettrix Technology; Reiser and Dickinson, 2008), in which the flight behavior was monitored using two microphones, each recording the sound of the left or right wingbeat. Sound data were analyzed in real-time to detect turning behavior and set the virtual heading direction. Visual and olfactory stimuli were updated in closed-loop based on the fly’s heading direction. To optogenetically silence DANs through the activation of GtACR, the end of an optic fiber (M118L03, Thorlabs) coupled to 530 nm LED (M530F2 Thorlabs) was placed 5∼10 mm above the fly, aiming at the head.

Flies were raised on a cornmeal agar with or without ATR for more than two days before the experiment. One day prior to the experiment, the cuticle above the fly’s head was removed and sealed with transparent, ultraviolet-curing adhesive (NOA 63, Norland) under cold anesthesia to increase the efficiency of optogenetic stimulation. Immediately before the experiment, flies starved for 4-7 h with water were cold anesthetized, tethered to a stainless-steel pin, and transferred to the flight simulator. Recording was initiated ∼3 min after the recovery from anesthesia.

Three odors with different values (γ-butyrolactone, neutral; 2-pentanone, attractive; 2-methylphenol, aversive) were each applied 3 times in a row with 4 s stimulation and 6 s inter-stimulus periods in a trial (Figure 6B). During the stimulation period, an odor was applied whenever the flies were in a restricted spatial region (45°) centered at the fly location at the time of odor contact. Air was applied outside of this region at the same flow rate. An experiment consisted of 7 trials and the order of odor presentation was randomized in each trial. For optogenetic suppression of DANs, LED light was applied only when the fly was within the odor. Experiment was terminated if the flies did not fly or turn much in the first several trials.

### Data Analysis

#### Analysis of DAN imaging data

Image processing and data analyses were conducted using custom codes written in MATLAB (Release 2018b, 2019a, 2021a, MathWorks). Ca^2+^ imaging data of DANs were analyzed in three steps (Figure 1C): (1) nonlinear registration of the template volume of the 15 MB compartments (Aso et al., 2014a) to MB247-DsRed signals in a master image of individual brains using ANTs (Avants et al., 2010, 2011), (2) correction for brain motion by rigid registration of time lapse images to the master image using GCaMP6f signals based on cross correlation, and (3) extraction of fluorescence changes in each compartment using the registered template.

Normalized changes in fluorescence (ΔF/F, (F – F_baseline_)/F_baseline_) were calculated using baseline fluorescence (F_baseline_) averaged over 3 s preceding the stimulus onset. Stimulus responses were reported as an average over 5 time frames covering the odor stimulation period unless otherwise noted. Stimulus onset was defined as the time when odors actually reach the fly, which was measured in a previous study (Badel et al., 2016).

Principal component analysis was performed with odor responses averaged across 10-11 animals. Odor value was decoded with partial least-squares regression and cross-validation was used to control for overfitting. The model was trained with data corresponding to all but one odor, and the value of the held-out odor was predicted. This procedure was repeated for all the training and test odor combinations. The accuracy of prediction was assessed using the coefficient of determination. To compare the information coding in DANs and PNs, we used PN responses to the same odors obtained in Badel et al. (2016). For classification of odor identity, odor responses in three out of four blocks were used to train the linear decoder and the remaining responses were used as test data. The classification accuracy was assessed by the percentage of correct output.

#### Analysis of MBON imaging data

To correct for brain movement, time lapse images were aligned to the first image using rigid transformation. Because spilt Gal4 lines were used to label single types of MBONs, the entire volume mostly covering the dendrites were used to calculate ΔF/F.

#### Connectome analysis

Connectome data was analyzed in R (version 4.0.5, R studio version 1.4.1106). The connectome data was downloaded from neuPRINT+ version v1.2.1 (https://neuprint.janelia.org/) using custom query. For example, the following query can be used to extract the pathways between PNs and PAM DANs transiting two interneurons via synapses stronger than weight 5, and obtain the identity of interneurons as well as the synaptic weights between neurons.

MATCH (i:Neuron)-[w1:ConnectsTo]->(h1:Neuron)-[w2:ConnectsTo]->(h2:Neuron)-[w3:ConnectsTo]->(o:Neuron) WHERE i.type CONTAINS “PN” AND o.type CONTAINS “PAM” AND w1.weight>5 AND w2.weight>5 AND w3.weight>5 RETURN i.bodyId, i.type, w1.weight, h1.type, h1.bodyId, w2.weight, h2.type, h2.bodyId, w3.weight, o.bodyId, o.type

All the pathways between PNs and DANs transiting one or two interneurons via synapses stronger than weight 5 were extracted, and a neural network was built with all the neurons contributing to these pathways. Interneurons were classified into three groups; group 1 receives input from PNs and other interneurons and sends output to other interneurons or DANs; group 2 receives input from PNs and sends output to other interneurons; group 3 receives input from other interneurons and sends output to DANs. The network was used to predict the odor responses of DANs from those of PNs recorded previously (Badel et al., 2016). PNs in the same glomerulus were given the same activity because they transmit highly correlated information (Kazama and Wilson, 2009). The output of neuron *j* was calculated from the activity of presynaptic neurons as follows;

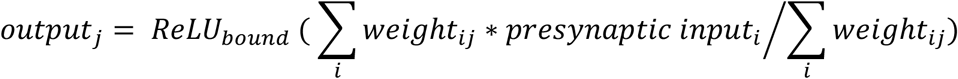

where

*ReLU*_*bound*_: rectified linear unit with an upper bound (set to 100)

*weight*_*ij*_: synaptic weight between input neuron *i* and output neuron *j*

*presynaptic input*_*i*_: odor response of presynaptic neuron *i* and the sum is taken over all the presynaptic neurons *i* connected to the output neuron *j*. The sign of synaptic weights for each neuron was set according to the metadata in hemibrain version v1.2.1 (Clements et al. 2020). Cholinergic neurons were set to excitatory and GABAergic and glutamatergic neurons were set to inhibitory. The number of neurons communicating with excitatory, inhibitory, or undetermined sign in each group is shown in Table S3. To reflect the contribution of recurrent connections, the activity of neurons in group 1 was first calculated using input from PNs and neurons in group 2, before including the recurrent input from neurons within group 1. This procedure represents the first-order approximation of the processes that take place in a recurrent network formed within group 1. In the final step, the activity of 307 DANs is summed within each neuron type and then within each compartment to obtain the activity in 15 compartments.

**Table S3.** Transmitter profile by the interneuron groups.

|  | Group 1 | Group 2 | Group 3 |
| --- | --- | --- | --- |
| **Excitatory** | 1501 | 636 | 303 |
| **Inhibitory** | 15 | 291 | 76 |
| **Undetermined** | 27 | 411 | 589 |

#### Behavioral Data Analysis

Flight trajectories of the fly were calculated based on the absolute difference in the standardized left and right wing-beat amplitudes as described in Badel et al. (2016). The proportion of time that flies have spent outside of an odor region was calculated every 5 ms, and its average over the last 1 s of odor application period and over 7 trials was reported in Figures 6 and S7. The mean and SEM were computed across all the tested flies.

#### Statistical Analysis

Details of statistical analyses, including the statistical tests performed, significance, sample size, and dispersion, are reported either in the main text, figures, the corresponding figure legends, or Table S4. Sample sizes were predetermined based on the effect size and the variability of data observed in pilot experiments. None of the calcium imaging data were excluded. For behavioral data, trials were excluded from analyses when flies were spinning (turning more than 1440 deg/s) or not flying.

**Table S4.** Results of statistical analyses

| **Two way ANOVA followed by Tukey's multiple comparison test.** |  |  |  |  |  |
| --- | --- | --- | --- | --- | --- |
| **Figure 5C (aversive vs attractive odor, saline)** |  |  |  |  |  |
|  | **Sum of the squares** | **Degrees of freedom** | **Mean squared error** | **F-statistic** | **p-value** |
| **Repeat** | 4.38E+04 | 3 | 1.46E+04 | 3.534 | 0.0256 |
| **Odor** | 1.43E+05 | 1 | 1.43E+05 | 34.5651 | 1.54E-06 |
| **Interaction** | 877.4039 | 3 | 292.468 | 0.0707 | 0.9752 |
| **Error** | 1.32E+05 | 32 | 4.14E+03 |  |  |
| **Total** | 3.20E+05 | 39 |  |  |  |
| **Comparison** |  | **Lower 95% CI** | **Difference between estimated group means** | **Upper 95% CI** | **p-value** |
| **aversive** | **attractive** | 78.1341 | 119.5559 | 160.9777 | 1.54E-06 |
| **Figure 5D (aversive odor)** |  |  |  |  |  |
|  | **Sum of the squares** | **Degrees of freedom** | **Mean squared error** | **F-statistic** | **p-value** |
| **Repeat** | 9.60E+04 | 3 | 3.20E+04 | 6.5668 | 8.25E-04 |
| **Blocker** | 2.81E+04 | 2 | 1.40E+04 | 2.8821 | 0.0658 |
| **Interaction** | 6.18E+03 | 6 | 1.03E+03 | 0.2112 | 0.9716 |
| **Error** | 2.34E+05 | 48 | 4.87E+03 |  |  |
| **Total** | 3.64E+05 | 59 |  |  |  |
| **Comparison** |  | **Lower 95% CI** | **Difference between estimated group means** | **Upper 95% CI** | **p-value** |
| **Saline** | **SCH** | -88.2707 | -34.8745 | 18.5217 | 0.2642 |
| **Saline** | **Wash** | -36.2624 | 17.1338 | 70.53 | 0.7194 |
| **SCH** | **Wash** | -1.388 | 52.0083 | 105.4045 | 0.0577 |
| **Figure 5D (attractive odor)** |  |  |  |  |  |
|  | **Sum of the squares** | **Degrees of freedom** | **Mean squared error** | **F-statistic** | **p-value** |
| **Repeat** | 3.42E+04 | 3 | 1.14E+04 | 1.8438 | 0.1518 |
| **Blocker** | 2.46E+05 | 2 | 1.23E+05 | 19.8915 | 5.10E-07 |
| **Interaction** | 2.84E+03 | 6 | 474.0817 | 0.0766 | 0.9981 |
| **Error** | 2.97E+05 | 48 | 6.19E+03 |  |  |
| **Total** | 5.81E+05 | 59 |  |  |  |
| **Comparison** |  | **Lower 95% CI** | **Difference between estimated group means** | **Upper 95% CI** | **p-value** |
| **Saline** | **SCH** | -210.0799 | -149.9001 | -89.7202 | 6.85E-07 |
| **Saline** | **Wash** | -94.8556 | -34.6757 | 25.5041 | 0.3523 |
| **SCH** | **Wash** | 55.0445 | 115.2243 | 175.4041 | 8.20E-05 |
| **Figure 5G (aversive vs attractive odor, saline)** |  |  |  |  |  |
|  | **Sum of the squares** | **Degrees of freedom** | **Mean squared error** | **F-statistic** | **p-value** |
| **Repeat** | 0.6816 | 3 | 0.2272 | 7.0568 | 6.45E-04 |
| **Odor** | 0.3261 | 1 | 0.3261 | 10.1285 | 0.0028 |
| **Interaction** | 0.0502 | 3 | 0.0167 | 0.5193 | 0.6714 |
| **Error** | 1.2879 | 40 | 0.0322 |  |  |
| **Total** | 2.3458 | 47 |  |  |  |
| **Comparison** |  | **Lower 95% CI** | **Difference between estimated group means** | **Upper 95% CI** | **p-value** |
| **aversive** | **attractive** | -0.2695 | -0.1649 | -0.0602 | 0.0028 |
| **Figure 5H (aversive odor)** |  |  |  |  |  |
|  | **Sum of the squares** | **Degrees of freedom** | **Mean squared error** | **F-statistic** | **p-value** |
| **Repeat** | 0.4675 | 3 | 0.1558 | 1.464 | 0.2334 |
| **Blocker** | 4.4944 | 2 | 2.2472 | 21.1113 | 1.14E-07 |
| **Interaction** | 0.1344 | 6 | 0.0224 | 0.2104 | 0.9722 |
| **Error** | 6.3867 | 60 | 0.1064 |  |  |
| **Total** | 11.4829 | 71 |  |  |  |
| **Comparison** |  | **Lower 95% CI** | **Difference between estimated group means** | **Upper 95% CI** | **p-value** |
| **Saline** | **SCH** | 0.3569 | 0.5832 | 0.8096 | 1.74E-07 |
| **Saline** | **Wash** | -0.0953 | 0.1311 | 0.3574 | 0.3516 |
| **SCH** | **Wash** | -0.6785 | -0.4522 | -0.2258 | 3.21E-05 |

**Figure 5H (attractive odor)**
|  | **Sum of the squares** | **Degrees of freedom** | **Mean squared error** | **F-statistic** | **p-value** |
| --- | --- | --- | --- | --- | --- |
| **Repeat** | 1.8325 | 3 | 0.6108 | 7.1182 | 3.61E-04 |
| **Blocker** | 0.5128 | 2 | 0.2564 | 2.9879 | 0.0579 |
| **Interaction** | 0.2264 | 6 | 0.0377 | 0.4398 | 0.8493 |
| **Error** | 5.1488 | 60 | 0.0858 |  |  |
| **Total** | 7.7206 | 71 |  |  |  |
| **Comparison** |  | **Lower 95% CI** | **Difference between estimated group means** | **Upper 95% CI** | **p-value** |
| **Saline** | **SCH** | -0.3948 | -0.1916 | 0.0116 | 0.0685 |
| **Saline** | **Wash** | -0.2319 | -0.0286 | 0.1746 | 0.9388 |
| **SCH** | **Wash** | -0.0403 | 0.163 | 0.3662 | 0.1399 |

**Figure 6E (aversive odor)**
|  | **Sum of the squares** | **Degrees of freedom** | **Mean squared error** | **F-statistic** | **p-value** |
| --- | --- | --- | --- | --- | --- |
| **Group** | 1.5925 | 3 | 0.5308 | 4.5781 | 0.0039 |
| **Repeat** | 2.7433 | 2 | 1.3717 | 11.8293 | 1.25E-05 |
| **Interaction** | 0.466 | 6 | 0.0777 | 0.6697 | 0.6742 |
| **Error** | 28.2929 | 244 | 0.116 |  |  |
| **Total** | 33.094 | 255 |  |  |  |
| **Comparison** |  | **Lower 95% CI** | **Difference between estimated group means** | **Upper 95% CI** | **p-value** |
| **ATR-LED+** | **ATR-LED-** | -0.144 | 0.0076 | 0.1591 | 0.9993 |
| **ATR-LED+** | **ATR+LED+** | -0.2686 | -0.117 | 0.0346 | 0.1946 |
| **ATR-LED+** | **ATR+LED-** | -0.0418 | 0.1111 | 0.2641 | 0.2423 |
| **ATR-LED-** | **ATR+LED+** | -0.2817 | -0.1245 | 0.0326 | 0.1749 |
| **ATR-LED-** | **ATR+LED-** | -0.0549 | 0.1036 | 0.262 | 0.3345 |
| **ATR+LED+** | **ATR+LED-** | 0.0697 | 0.2281 | 0.3866 | 0.0012 |

**Figure 6E (neutral odor)**
|  | **Sum of the squares** | **Degrees of freedom** | **Mean squared error** | **F-statistic** | **p-value** |
| --- | --- | --- | --- | --- | --- |
| **Group** | 2.1634 | 3 | 0.7211 | 10.81 | 1.08E-06 |

| **Repeat** | 0.5872 | 2 | 0.2936 | 4.4015 | 0.0132 |
| --- | --- | --- | --- | --- | --- |
| **Interaction** | 0.2179 | 6 | 0.0363 | 0.5444 | 0.7741 |
| **Error** | 16.277 | 244 | 0.0667 |  |  |
| **Total** | 19.2807 | 255 |  |  |  |
| Comparison |  | Lower 95% CI | Difference between estimated group means | Upper 95% CI | p-value |
| **ATR-LED+** | **ATR-LED-** | -0.1026 | 0.013 | 0.1285 | 0.9916 |
| **ATR-LED+** | **ATR+LED+** | -0.2524 | -0.1379 | -0.0234 | 0.0106 |
| **ATR-LED+** | **ATR+LED-** | 0.0102 | 0.1262 | 0.2422 | 0.0267 |
| **ATR-LED-** | **ATR+LED+** | -0.2701 | -0.1509 | -0.0316 | 0.0063 |
| **ATR-LED-** | **ATR+LED-** | -0.0075 | 0.1132 | 0.2339 | 0.0753 |
| **ATR+LED+** | **ATR+LED-** | 0.1444 | 0.2641 | 0.3838 | 8.60E-08 |

**Figure 6E (attractive odor)**
|  | Sum of the squares | Degrees of freedom | Mean squared error | F-statistic | p-value |
| --- | --- | --- | --- | --- | --- |
| **Group** | 1.4576 | 3 | 0.4859 | 10.7636 | 1.21E-06 |
| **Repeat** | 0.5773 | 2 | 0.2887 | 6.3946 | 0.002 |
| **Interaction** | 0.225 | 6 | 0.0375 | 0.8308 | 0.5471 |
| **Error** | 10.3369 | 229 | 0.0451 |  |  |
| **Total** | 12.6819 | 240 |  |  |  |
| Comparison |  | Lower 95% CI | Difference between estimated group means | Upper 95% CI | p-value |
| **ATR-LED+** | **ATR-LED-** | -0.0479 | 0.0508 | 0.1496 | 0.5486 |
| **ATR-LED+** | **ATR+LED+** | -0.2063 | -0.1105 | -0.0147 | 0.016 |
| **ATR-LED+** | **ATR+LED-** | 0.0062 | 0.1038 | 0.2014 | 0.0319 |
| **ATR-LED-** | **ATR+LED+** | -0.2639 | -0.1613 | -0.0587 | 3.12E-04 |
| **ATR-LED-** | **ATR+LED-** | -0.0513 | 0.053 | 0.1572 | 0.56 |
| **ATR+LED+** | **ATR+LED-** | 0.1128 | 0.2143 | 0.3158 | 3.46E-07 |

## Supplemental Figures

**Figure S1.**
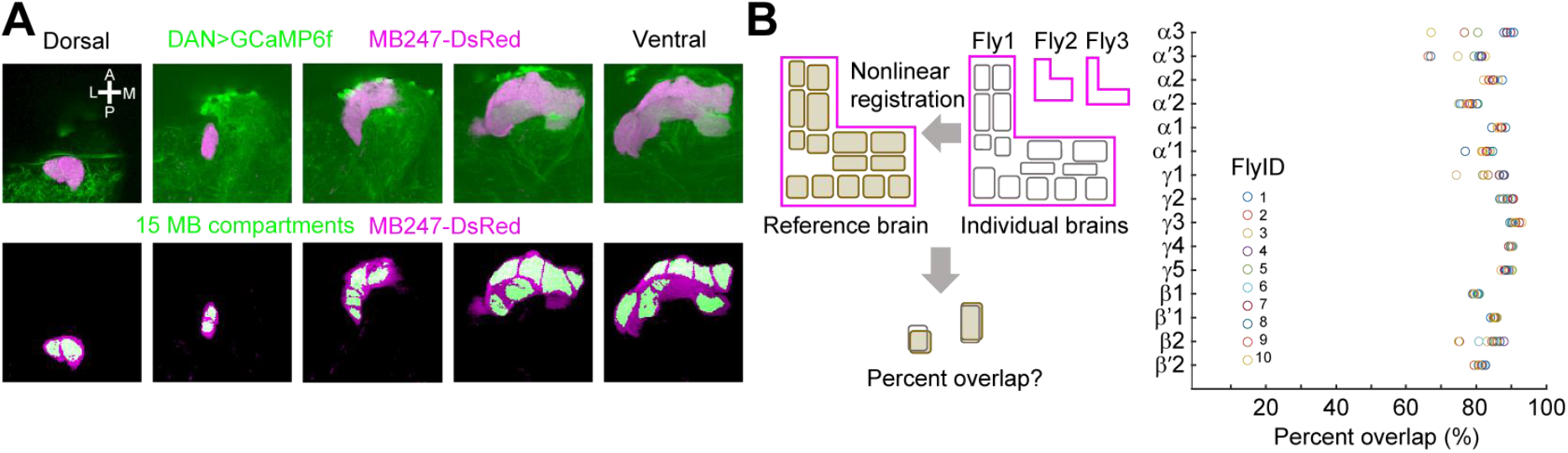
Assessment of the quality of the registration (A) Schematic of the template MB compartments registered to an example brain. (Top) GCaMP6f signals (green) and MB247-DsRed signals (magenta) at representative z-planes along dorsoventral axis. (Bottom) Registered 15 MB compartments (green) overlaid on MB247-DsRed signals in the corresponding z-planes. (B) The quality of the registration was assessed by computing, for each compartment, the percent overlap in volume between the same type of compartments registered to different brains. This comparison was repeated 10 times, each time with a different brain. The average percent overlap is reported for each compartment.

**Figure S2.**
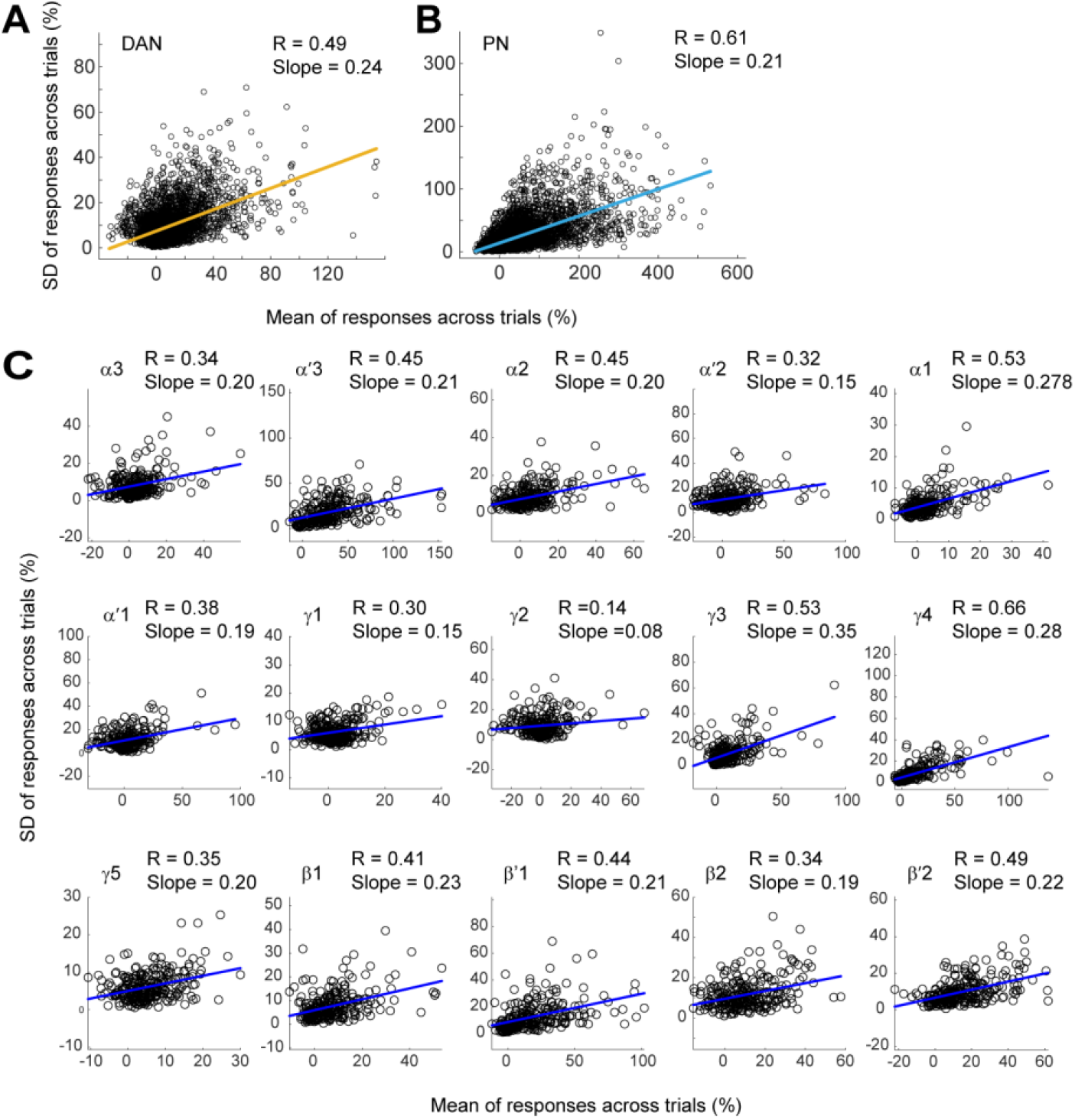
Odor responses in DANs and PNs show similar trial-to-trial variability (A, B) Trial-to-trial variability of odor responses of DANs in 15 compartments (A) and PNs in 37 glomeruli (B). Scatter plots show the mean and SD of responses across trials, calculated for each of the 25 odors. Data from individual flies (10–11 flies for DANs and 4–9 flies for PNs) are pooled. DANs and PNs show similar normalized response variability captured by the slope. (C) Trial-to-trial variability of odor responses of DANs in individual compartments. Same data as in A.

**Figure S3.**
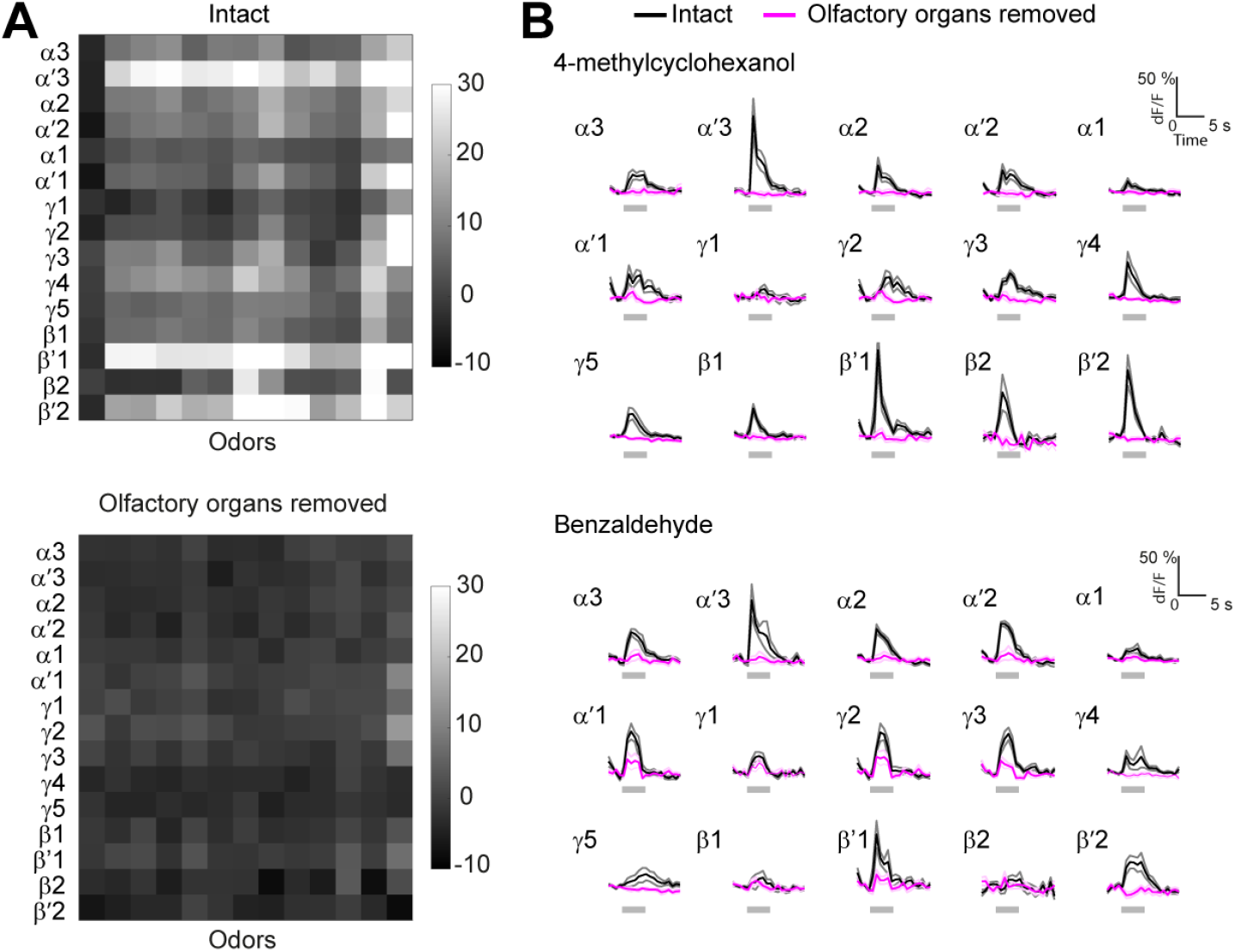
Odor-evoked DAN responses mostly originate in olfactory organs (A) Average peak responses of DANs to 13 odors with or without antennae and maxillary palps. Average is taken across time (odor application period), trials (2 trials per condition), and flies (n = 5 flies). Odor identity from the left; air, 2-pentanone, ethyl butyrate, 2-methylphenol, apple cider vinegar, ethyl acetate, 3-methylthio-1-propanol, 3-octanol, propionic acid, mineral oil, water, 4-methylcyclohexanol, and benzaldehyde. (B) Comparison between responses of DANs to two odors before (black) and after (magenta) the removal of olfactory organs (n = 5 flies, average across trials and flies, same data as in A).

**Figure S4.**
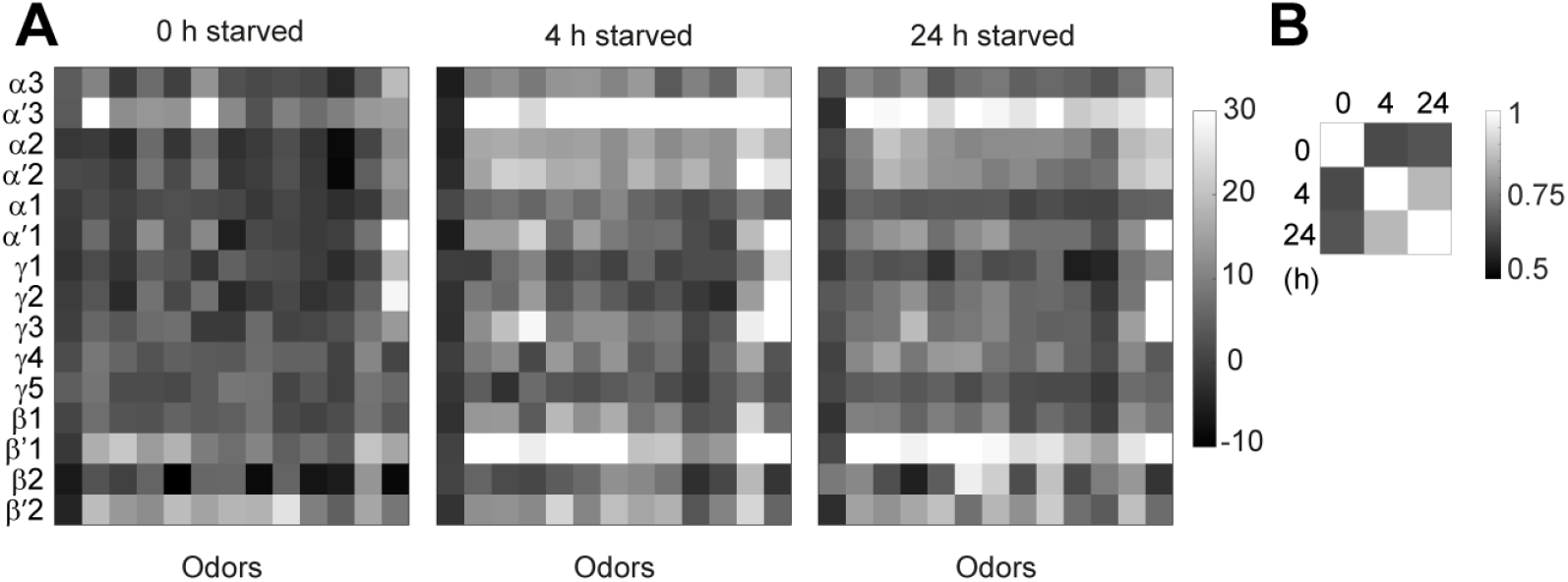
Odor-evoked DAN responses in fed and starved conditions (A) Average peak responses of DANs to 13 odors under three different feeding conditions (average across odor application period, 4 trials, and flies, n = 5, 10, and 5 flies for 0, 4, and 24 h starved conditions). Odor identity from the left; air, 2-pentanone, ethyl butyrate, 2-methylphenol, apple cider vinegar, ethyl acetate, 3-methylthio-1-propanol, 3-octanol, propionic acid, mineral oil, water, 4-methylcyclohexanol, and benzaldehyde. (B) Pearson’s correlation between response matrices in A.

**Figure S5.**
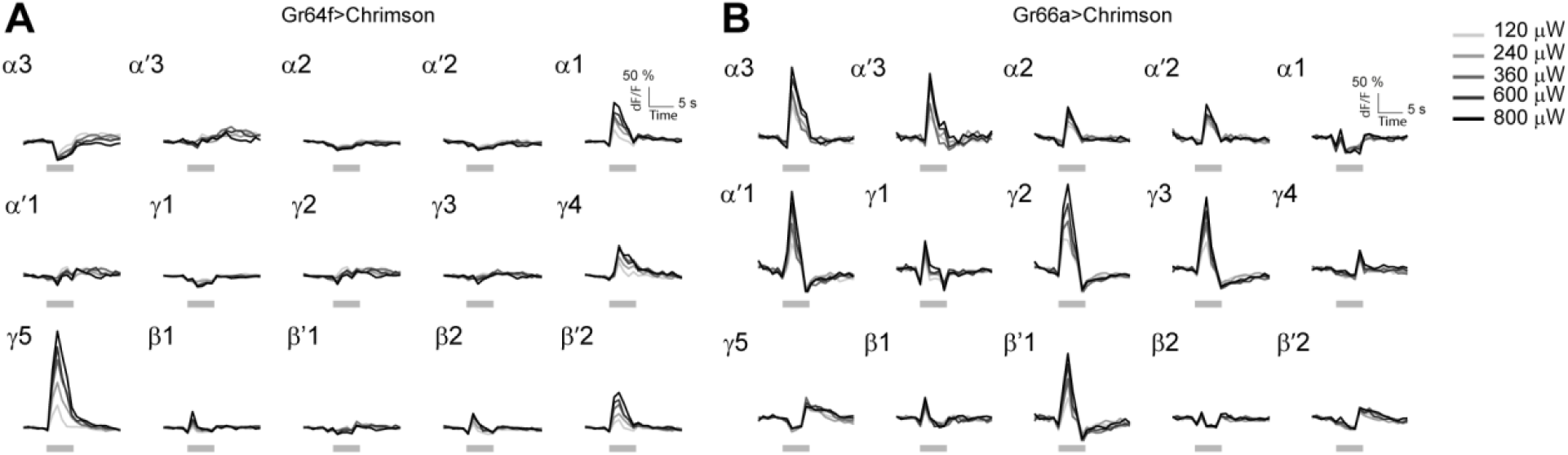
DAN responses to various levels of optogenetic GRN stimulation (A) Average responses of DANs to optogenetic stimulation of Chrimson-expressing Gr64f GRNs (average across 4 trials and 5 flies). Stimulation with increasing light intensity evoked increasing responses. 240 μW was used to examine the integration of gustatory and olfactory inputs by DANs in Figure 4. (B) Same as in A but for optogenetic stimulation of Chrimson-expressing Gr66a GRNs (n = 5 flies).

**Figure S6.**
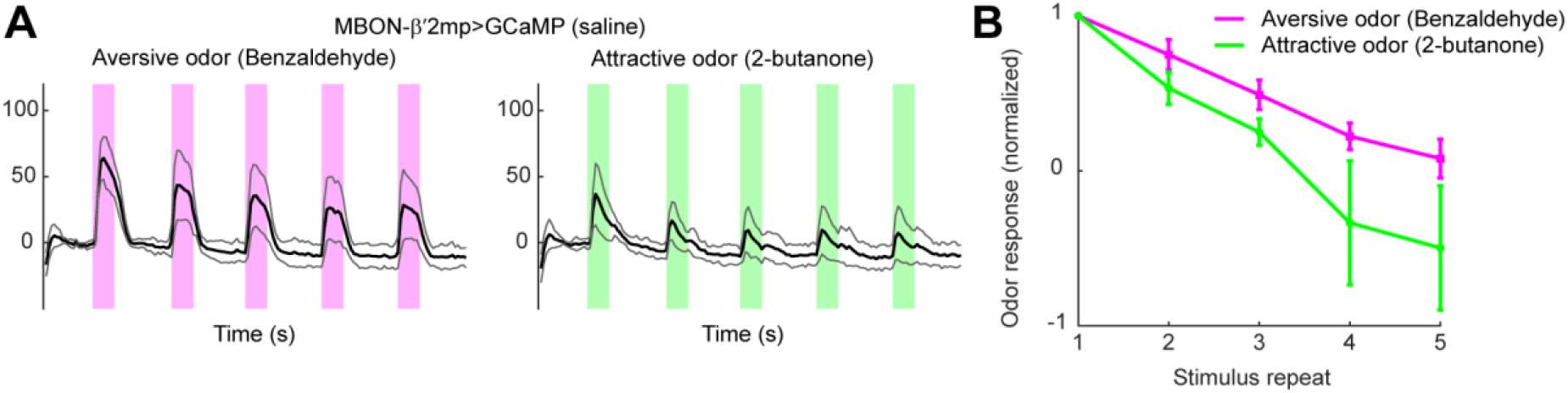
Odor value-dependent plasticity is observed in MBON-β’2mp also with a different odor pair (A) Responses of MBON-β’2mp to repetitive application of aversive (Benzaldehyde) and attractive (2-butanone) odors (n = 5 flies). (B) Responses are more depressed in response to an attractive over aversive odor (p = 0.01, two-way ANOVA). Error bar represents SEM.

**Figure S7.**
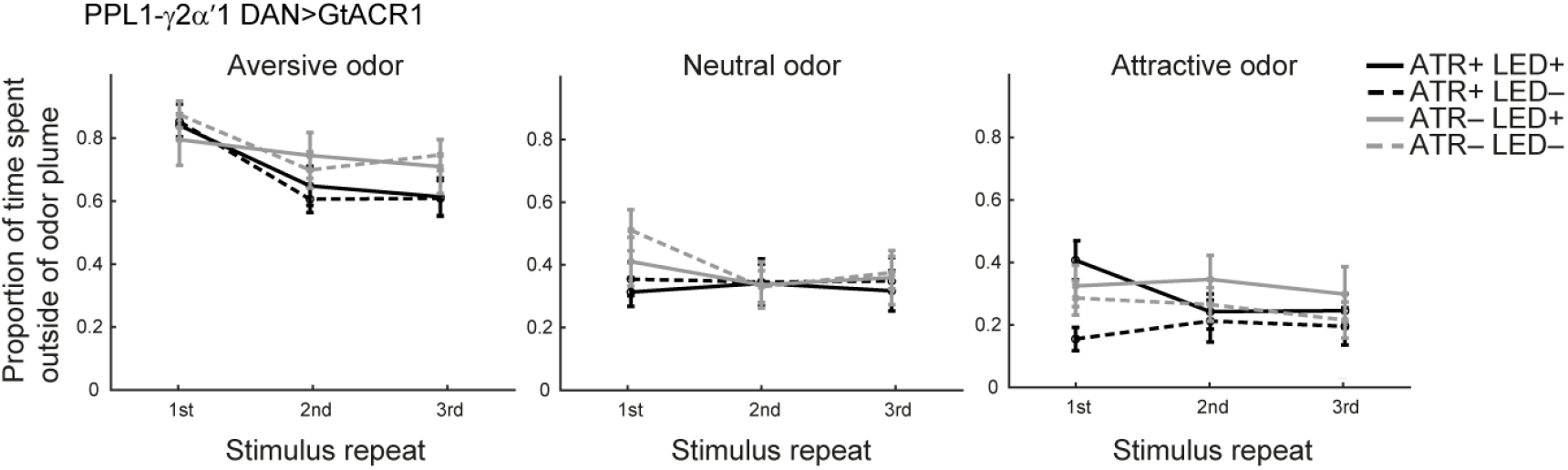
Behavioral effect of suppressing PPL1-γ2α’1 DAN activity Same as in Figure 6E, but for suppressing the activity of PPL1-γ2α’1 DANs (n = 17–23 flies for each condition). Optogenetic suppression had little effect, presumably reflecting a complex action of dopamine on MBONs in the corresponding γ2/α’1 compartment (Figure 5F-H, p = 0.29, 0.50, and 0.057, for aversive, neutral, and attractive odors, two-way ANOVA).

